# Lateral entorhinal cortex suppresses drift in cortical memory representations

**DOI:** 10.1101/2021.11.04.467279

**Authors:** Maryna Pilkiw, Justin Jarovi, Kaori Takehara-Nishiuchi

## Abstract

Memory retrieval is thought to depend on the reinstatement of cortical memory representations guided by pattern completion processes in the hippocampus. The lateral entorhinal cortex (LEC) is one of the intermediary regions supporting hippocampal-cortical interactions and houses neurons that prospectively signal past events in a familiar environment. To investigate the functional relevance of the LEC’s activity for cortical reinstatement, we pharmacologically inhibited the LEC and examined its impact on the stability of ensemble firing patterns in one of the LEC’s efferent targets, the medial prefrontal cortex (mPFC). When male rats underwent multiple epochs of identical stimulus sequences in the same environment, the mPFC maintained a stable ensemble firing pattern across repetitions, particularly when the sequence included pairings of neutral and aversive stimuli. With LEC inhibition, the mPFC still formed an ensemble pattern that accurately captured stimuli and their associations within each epoch. However, LEC inhibition markedly disrupted its consistency across the epochs by decreasing the proportion of mPFC neurons that stably maintained firing selectivity for stimulus associations. Thus, the LEC stabilizes cortical representations of learned stimulus associations, thereby facilitating the recovery of the original memory trace without generating a new, redundant trace for familiar experiences. Failure of this process might underlie retrieval deficits in conditions associated with degeneration of the LEC, such as normal aging and Alzheimer’s disease.

**SIGNIFICANCE STATEMENT:** To recall past events, the brain needs to reactivate the activity patterns that occurred during those events. However, such reinstatement of memory traces is not trivial because it goes against the brain’s natural tendency to restructure the activity patterns continuously. We found that dysfunction of a brain region called the lateral entorhinal cortex (LEC) worsened the drift of the brain activity when rats repeatedly underwent the same events in the same room and made them behave as if they had never experienced these events before. Thus, the LEC stabilizes the brain activity to facilitate the recovery of the original memory trace. Failure of this process might underlie memory problems in elderly and Alzheimer’s disease patients with the degenerated LEC.

## INTRODUCTION

Memory retrieval is presumed to involve the reinstatement of cortical memory representations guided by the hippocampus, which pattern-completes the full content of the original experience from a partial cue (Teyler and DiScenna, 1986; Alvarez and Squire, 1994; McClelland et al., 1995; Teyler and Rudy, 2007). This long-standing view has received empirical support from recent human imaging (Ritchey et al., 2013; Staresina et al., 2013; Bosch et al., 2014; Horner et al., 2015) and rodent gene-based activity mapping studies (Tanaka et al., 2014; Guo et al., 2018; Guskjolen et al., 2018). In contrast, parallel studies tracked firing patterns of hippocampal neurons longitudinally and uncovered their major reorganization across repeated experiences (Mankin et al., 2012; Schimanski et al., 2013; Ziv et al., 2013; Hainmueller and Bartos, 2018; Taxidis et al., 2020). Although such representational drift may be instrumental to flexible memory encoding/updating (Mau et al., 2020), it would work against the stability required for memory retrieval, indicating that the hippocampus may not be the only region that coordinates cortical reinstatement (Frankland and Bontempi, 2005; Cowansage et al., 2014; Takehara-Nishiuchi, 2014; Goode et al., 2020).

The entorhinal cortex serves as an interface between the hippocampus and neocortex (Squire, 1992; Eichenbaum, 2000) and consists of the medial and lateral areas with dissociable anatomical connectivity and neural selectivity (Knierim et al., 2014; Nilssen et al., 2019). Neurons in the lateral entorhinal cortex (LEC) respond to physical stimuli that are currently present (Young et al., 1997; Deshmukh and Knierim, 2011; Xu and Wilson, 2012; Tsao et al., 2013; Igarashi et al., 2014; Keene et al., 2016; Pilkiw et al., 2017; Suter et al., 2019; Woods et al., 2020) and track their consistency across days (Deshmukh and Knierim, 2011; Tsao et al., 2013). Also, nearly all LEC neurons abruptly change baseline firing rates upon an entry to a familiar environment in a manner depending on which sensory events had taken place in the environment (Pilkiw et al., 2017). These findings raise the possibility that the LEC prospectively signals past events from cues in familiar environments, thereby facilitating the recovery of the corresponding cortical representations.

To test this idea, we investigated the impact of pharmacological inhibition of the LEC on the stability of memory-selective ensemble firing patterns in one of its efferent targets, the prelimbic region of the medial prefrontal cortex (PrL; (Swanson and Köhler, 1986; Insausti et al., 1997; Kerr et al., 2007; Ährlund-Richter et al., 2019). To create conditions of constant sensory experiences, we repeated multiple epochs of trace eyeblink conditioning (Woodruff-Pak and Disterhoft, 2008; Takehara-Nishiuchi, 2018) in the same environment during a daily recording session. This non-spatial associative learning paradigm is ideal for testing our hypothesis because its memory retrieval depends on the hippocampus (Kim et al., 1995; Takehara et al., 2003), LEC (Morrissey et al., 2012), and PrL (Takehara et al., 2003; Takehara-Nishiuchi et al., 2006). Moreover, the PrL houses neurons coding stimulus associations (Takehara-Nishiuchi and McNaughton, 2008; Hattori et al., 2014; Morrissey et al., 2017), and learning is accompanied by increased oscillatory coupling between the PrL and LEC (Takehara-Nishiuchi et al., 2011). We found that LEC inhibition exacerbated drift in PrL neuronal representations over time, thereby preventing accurate reinstatement of cortical memory traces of the learned stimulus associations.

## MATERIALS AND METHODS

### Animals

All surgical and experimental procedures were approved by the Animal Care and Use Committee at the University of Toronto (AUP 20011400). Four male Long-Evans rats (Charles River Laboratories, St. Constant, QC, Canada) weighing 450–600 g at the time of surgery were used in the experiment. The rats were single-housed in Plexiglas cages in a colony room with *ad libitum* access to food and water. They were maintained on a reversed 12-hour light-dark circadian cycle, and all experiments took place during the dark part of the cycle. Prior to the experiment, rats were handled 1–2 times per week.

### Electrodes for single-unit recording

Microdrives and electrodes were constructed in-house following a procedure described previously (Kloosterman et al., 2009). A 3D-printed plastic base housed twelve individually movable tetrodes, two reference electrodes, and an electrode interface board (EIB-54-Kopf, Neuralynx, Bozeman, MT, USA). Tetrodes were made by folding in four and twisting a 36 cm wire (12 µm polyimide coated nichrome wire, Sandvik, Stockholm, Sweden) following the procedure used in our previous work (Takehara-Nishiuchi and McNaughton, 2008; Morrissey et al., 2017). Each tetrode was connected to the plastic base with a shuttle containing a custom-made screw which allowed precise control of the tetrode movement. From the top, individual tetrode wires were connected to the interface board with gold pins. From the bottom, the electrode tips were cut and gold plated to reduce impedance to 200-250 kΩ. At the base, the tetrodes were arranged in a 7×2 bundle (∼2×0.5 mm).

### Surgical procedures

The electrode array was implanted above the prelimbic region of the medial prefrontal cortex with the same procedure as those used in our previous work (Takehara-Nishiuchi and McNaughton, 2008; Morrissey et al., 2017). All surgeries were conducted under aseptic conditions in a dedicated surgical suite. Rats were anesthetized with isoflurane (induced at 5% and maintained at 2–2.5% by volume in oxygen at a flow rate of 0.9-1.0 L/min; Halocarbon Laboratories, River Edge, NJ, United States) and placed in a stereotaxic holder with the skull surface in the horizontal plane. A craniotomy was opened between 3.3–5.4 mm anterior and 0.65–0.95 mm lateral to bregma. After removing the dura mater, the array was placed at the brain surface (1.5-1.75 mm ventral from the skull surface) at a 15° forward angle. In addition, microinfusion cannulae were implanted bilaterally in the lateral entorhinal cortex (6.45 mm posterior, 6.7–6.8 mm lateral and 8.4-8.9 mm ventral to Bregma). The craniotomy was covered with Gelfoam (Pharmacia & Upjohn, NJ, United States), and the array was held in place with self-adhesive resin cement (3M, MN, United States) and self-curing dental acrylic (Lang Dental Manufacturing, Wheeling, IL, United States), supported by screws inserted in the skull. Two additional ground screws were placed above the parietal cortex. During the surgery, all tetrodes were lowered 1.0–1.5 mm into the brain. For the next seven days, the rat was connected to the recording system daily to monitor the quality of neural signal and to adjust the location of tetrodes. During this period, each tetrode was gradually lowered (0.03–1 mm per day) to target tetrode tips to the prelimbic region at 2.0–3.6 mm ventral from the brain surface. Two reference electrodes remained superficially in the cortex (1 mm below the brain surface).

### Behavioral paradigm and pharmacological manipulations

The experimental apparatus consisted of a 50×50×50 cm black box fitted with a sliding door connected to a small hallway and a 6×12 cm perforated Plexiglas window. A speaker and a 5 mm LED were attached outside of the window. The conditioning began at least seven days after the surgery. All rats initially underwent two stages of conditioning, Acquisition (7-9 days) and Test stages (>14 days). All sessions in both stages took place in the same conditioning chamber.

*Acquisition* During the first two days, rats were placed in the conditioning chamber without receiving any stimulus presentations. On the third day, the conditioning began. The conditioned stimulus (CS) was presented for 100 ms and consisted of an auditory stimulus (85 dB, 2.5 kHz pure tone) or a visual stimulus (white LED light blinking at 50 Hz). The unconditioned stimulus (US) was a 100 ms mild electrical shock near the eyelid (100 Hz square pulse, 0.3–2.0 mA), and the intensity was carefully monitored via an overhead camera and adjusted to ensure a proper eyeblink/head turn response (Morrissey et al., 2012; Volle et al., 2016). The timing of the CS and US presentation was controlled by a microcomputer (Arduino Mega, Arduino, Italy). The US was generated by a stimulus isolator (ISO-Flex, A.M.P.I., Jerusalem, Israel) and applied through a pair of stainless-steel wires chronically implanted in the eyelid. The CS presentations were separated by a random interval ranging from 20 to 40 seconds. The conditioned response (CR) was defined as anticipatory blinking responses that occurred before US onset and detected by recording an electromyogram (EMG) from the left upper eyelid.

Each session consisted of two epochs of trace eyeblink conditioning, each of which included 100 trials of CS presentations. The two epochs were separated by a 10 min rest period. Each epoch began with 20 presentations of the CS by itself (CS-alone block), followed by 80 trials in which the CS was paired with the US, separated by a stimulus-free interval of 500 ms (CS-US block). The same CS was used in the CS-alone and CS-US blocks within each epoch. The CS-alone block always proceeded the CS-US block, enabling the rats to learn that the temporal context signaled the change in CS-US contingency (Takehara-Nishiuchi and McNaughton, 2008; Morrissey et al., 2017; Pilkiw et al., 2017). To make the rats form two distinct associative memories, the first epoch used one of the two CS (*e.g*., auditory CS), and the second epoch used the other CS (*e.g*., visual CS). Before and after each epoch, the rat was placed in the hallway separating the conditioning chambers. Daily sessions continued until a rat expressed CRs in at least 60% of CS-US trials in one of the blocks.

*Test* After the Acquisition stage, the rats underwent Test sessions with microinfusion of either artificial cerebral spinal fluid (aCSF) or the GABA_A_ receptor agonist, muscimol, bilaterally into the LEC. This paper reports neuronal activity recorded during these sessions. In each daily session, the rats underwent four epochs, each of which started with a block of 20 CS-alone trials, followed by a block of 50 CS-US trials. Either tone or light CS was used throughout the four epochs on a given day. Before and after each epoch, the rats had a 5 min rest period in the hallway. At the end of the second epoch, the rat was moved to a small box and gently wrapped with a clean towel without being anesthetized. As in our previous work (Morrissey et al., 2012), 1 µl of a solution of muscimol hydrobromide (1 µg in aCSF, G019 Sigma-Aldrich, Canada), or aCSF (Tocris Bioscience, Canada) was infused through the chronically implanted cannula over two minutes. The infusion cannula was left in place for an additional minute before extraction. Then, the rat was moved to the hallway and allowed to rest for 30-60 min before the third epoch began. In each rat, the CS type (tone or light) and drug type (aCSF or muscimol) were randomly assigned to each session (Table 1). In the first two Test sessions, some rats (PLN7, 8, 11) underwent the infusion procedure without actually infusing the drug solution to habituate them to the procedure.

**Table 1.**
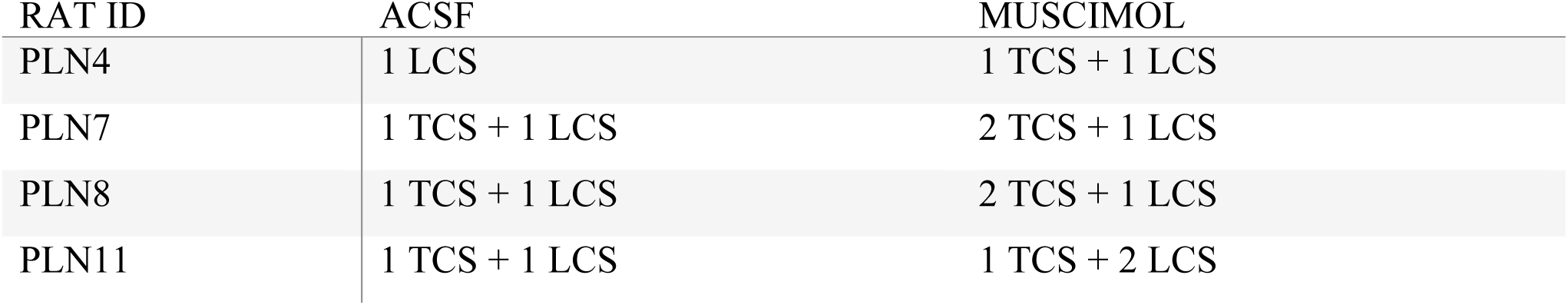
The number of sessions that each rat underwent with each drug and CS type. TCS, tone conditioned stimulus; LCS, light conditioned stimulus.

### Data acquisition

During the Test sessions, we simultaneously recorded action potentials from individual neurons in the right prelimbic cortex and EMG activity from the left upper eyelid. Action potentials were captured using the tetrode technique (Wilson and McNaughton, 1993). Rats were connected to the recording system through an Electrode Interface Board (EIB-54-Kopf, Neuralynx, Bozeman, MT, United States) contained within the microdrive array fixed to the animal’s head. The EIB was connected to a headstage (CerePlex-M 64, Blackrock Microsystems, UT, United States), and signals were acquired through the Cerebus Neural Signal Processor (Blackrock Microsystems, UT, United States). A threshold voltage was set at 40–75 mV, and if the voltage on any channel of a tetrode exceeded this threshold, activity was collected from all four channels of the tetrode. The spiking activity of single neurons was sampled for 1 ms at 30 kHz, and signals were amplified and filtered between 600–6,000 Hz. EMG activity was continuously sampled at 6,108 Hz and filtered between 300–3,000 Hz.

### Behavioral Analysis

Learning of the association between the CS and US was measured based on the frequency of adaptive conditioned responses (CR), which were defined as a significant increase in EMG amplitude before US onset (Morrissey et al., 2012, 2017). The instantaneous amplitude of the EMG signal was calculated as the absolute value of the Hilbert transform of the EMG signal. Then, two values were calculated in each trial: 1) the averaged amplitude of EMG signals during a 200 ms period before CS onset (pre-CS value) and 2) the averaged amplitude during a 200 ms period before US onset or the corresponding interval in the CS-alone trials (CR value). A threshold was defined as the average of all pre-CS values plus one standard deviation. If the pre-CS value was greater than 130% of the threshold in a trial, the trial was discarded due to the hyperactivity of the rat immediately before the CS presentation. If the pre-CS value was below the threshold and CR-value was greater than 110% of the threshold, the trial was considered to have a CR. Finally, in the case when the pre-CS value was between the threshold and 130% of the threshold, the trial was considered to have a CR only if the CR value above the threshold was five times greater than the pre-CS value above the threshold. CR% was calculated as a ratio of the number of trials containing the CR to the total number of valid trials (*i.e.*, total trials minus hyperactive trials). CR% was calculated separately in each block in each session. The values were then averaged across all test sessions in each rat.

### Neural activity analysis

*Pre-processing* Putative units were isolated offline using an automated clustering software package (KlustaKwik, K.D., Harris, Rutgers, The State University of New Jersey, Newark, NJ, USA), followed by manual sorting (MClust, D.A. Redish, University of Minnesota, Minneapolis, MN; Waveform Cutter, S.L. Cowen, University of Arizona, Tucson, AZ, USA). Subsequent data analyses used only units in which the amplitude and shape of the spike waveform were consistent across the entire recording period.

*Firing-rate matrix correlations* This analysis used all 20 CS-alone trials and the first 20 CS-US trials in each epoch to match the number of trials between the two trial types. From each of 300 randomly subsampled neurons, we first extracted the spiking activity during a 1.6-sec window that covered from 0.4 s before to 1.2 s after CS onset in each block in each epoch. The spikes were then binned into sixteen 100-msec bins and averaged across trials. The 10^th^ bin was removed from the matrix to avoid the contamination of US artifacts in CS-US trials. The averaged value was normalized by dividing the firing rate in each bin by the maximum among the bins within each trial block in each epoch. Correlation coefficients were calculated between the matrix of the normalized firing rates (300 neurons × 15 bins) between two adjacent epochs. The procedure was repeated 20 times, each of which included a different set of randomly selected 300 neurons.

*Neural trajectory analysis* From each neuron, we first extracted the spiking activity during a 2.6-sec window centering at CS onset in CS-US or CS-alone block in each epoch. The activity during US was removed to avoid the contamination of US artifacts in CS-US trials. The spikes were then binned into 300-msec bins with a 50-msec increment. The binned firing rate was concatenated across four epochs, converted into a z-score, and stored in the firing rate matrix. Principal component analysis (PCA) was performed on the firing rate matrix. The top three principal components were averaged across trials in each epoch and plotted to visualize ensemble activity in neural state space.

*Ensemble decoding analysis* As in our previous studies (Morrissey et al., 2017; Xing et al., 2020), we quantified the selectivity of ensemble firing patterns by using decoding accuracy and classification errors of support vector machine (SVM) classifiers. SVM classifiers produce a model from training data attributes and then predict the target values using only the test data attributes. In our case, the attributes were the normalized binned firing rates of neurons, and the target values were the task phases, including the pre-CS, CS, CS-US interval, and post-US periods. All algorithms were run in MATLAB using the freely accessible LIBSVM library that implements the one-against-one approach for multi-class classifications (Chang and Lin, 2011). In each of 300 randomly sub-sampled neurons, we calculated firing rates in four 300-msec windows, the pre-CS (-300 to 0 msec after CS onset), CS (0 to 300 msec after CS onset), CS-US interval (300 to 600 msec after CS onset), and post-US (0 to 300 msec after US offset). The binned firing rate was concatenated across neurons and stored in a firing rate matrix. “Within-epoch” decoding was used to quantify ensemble selectivity for the task phase in each epoch. In each neuron, the firing rate in each trial was divided by the maximum firing rate of the neuron across all trials in an epoch (*i.e.*, 20 CS-alone and 50 CS-US trials). The classifiers were trained with the radial basis function (RBF) kernels. Two parameters in the RBF kernel, cost (C) and gamma were first identified by performing a grid search in which five-fold cross-validation accuracy was compared across multiple SVM runs with a different combination of C (11 values) and gamma (10 values). Because the degree of over-fitting should be negatively correlated with the accuracy, we selected the set of C and gamma that resulted in the highest accuracy. These parameters were then used for the training of SVM classifiers with 15 randomly selected trials. The other 5 trials were used for testing. The training and testing were repeated five times, each of which used a different set of 15 trials for training and the remaining 5 trials for testing. Decoding accuracy was defined as the proportion of test trials that were classified correctly. The procedure was repeated 20 times, each of which used a different set of 300 randomly selected neurons. “Cross-epoch” decoding was used to quantify the stability of ensemble firing patterns across two adjacent epochs. In this case, all trials from the preceding epoch were used for the training, and all trials from the following epoch were used for the testing.

As an alternative approach for quantifying ensemble similarity, we also calculated Pearson correlation coefficients between two matrices of 300 neurons × 4 task phases. For the stability within each epoch, trials in each epoch were randomly split in half, and firing rates were averaged across these trials to construct two firing rate matrices. For stability across two epochs, firing rates were averaged across all trials in each epoch. This process was repeated 20 times, each of which used a different set of 300 randomly subsampled neurons. To estimate the chance-level similarity, one of the matrices was shuffled across neurons and task phases before calculating the Pearson correlation coefficient.

*Single-neuron firing rate analysis* To quantify the stability of firing rates across trials in each epoch, firing rates of each neuron were calculated in four 300-msec windows, the pre-CS (- 300 to 0 msec after CS onset), CS (0 to 300 msec after CS onset), CS-US interval (300 to 600 msec after CS onset), and post-US (0 to 300 msec after US offset) in each trial. Then these rates were averaged across a series of bins of five trials (20 CS-alone trials divided into 4 bins; 50 CS-US trials divided into 10 bins). We then calculated Kullback-Leibler divergence between the binned firing rates and a uniform distribution. This was done separately before (Epoch 2) and after (Epoch 3) the infusion, and the change in the measure from Epoch 2 to Epoch 3 was calculated.

To quantify the degree to which each neuron changed firing rates after the presentation of the CS or US, firing rates were averaged across trials during the CS (a 600-ms window from CS onset to US onset; FR_CS), US (a 300-ms window starting from 100 ms after US offset; FR_US), and Baseline (a 600-ms window starting from 1 second before CS onset; FR_Pre).

Then, the Response Index (RI) as

RI = (FR_Stim – FR_Pre) / (FR_Stim + FR_Pre) where FR_Stim was either FR_CS or FR_US.

To quantify the degree to which each neuron differentiated CS-evoked firing rates between the CS-US block and CS-alone, we calculated the differential index (DI) as

DI = (FR_CS in CS-US – FR_CS in CS-alone) / (FR_CS in CS-US + FR_CS in CS-alone).

In each neuron, random permutation tests were performed by reassigning the labels (*i.e.*, baseline or CS-evoked; CS-US or CS-alone) to the firing rate in randomly chosen trials and calculating the firing rate change in the same manner as that with the real labels (100 iterations). This generated the probability distribution of RI or DI at the chance level. If the actual RI or DI fell in the top or bottom 2.5% of this distribution, the neuron was considered as “stimulus-responding” or “association-selective”.

*CR versus No-CR analyses* To compare the ensemble stability between trials in which the rats expressed CRs (CR trials) and those in which they did not (No-CR trials), we first selected sessions in which the rats expressed CRs in at least 30% of trials (Epoch 3, 7 aCSF sessions, 8 muscimol sessions; Epoch 4, 7 aCSF sessions, 5 muscimol sessions). In each session, CR and No-CR trials were randomly split into two sets. Firing rates were calculated in four 300-msec windows, the pre-CS (-300 to 0 msec after CS onset), CS (0 to 300 msec after CS onset), CS-US interval (300 to 600 msec after CS onset), and post-US (0 to 300 msec after US offset) in each trial. Then these rates were averaged across the two sets of trials to construct two firing rate matrices. Pearson correlation coefficients were calculated between the two firing rate matrices.

### Statistical analysis

The data was presented as the group mean ± standard error of the mean or 95% confidence interval. Statistical analysis was performed with SPSS Statistics or MATLAB. To determine the statistical significance, we used two-way analysis of variance (ANOVA), t-tests, Wilcoxon signed-rank, rank-sum, or binominal tests. Significance was defined as \**P* < 0.05, \*\**P* < 0.01, \*\*\**P* < 0.001.

### Histology

Upon completion of all recordings, the location of electrodes was marked by electrolytic lesions. For each tetrode, 5 µA was passed through one wire of each tetrode (positive to the electrode, negative to the animal ground) for 20 s. After 3-5 days after lesioning, rats were perfused intracardially with 0.9% saline followed by 10% buffered formalin. The brain was removed from the skull and stored in 10% formalin for 1-2 days. For cryosectioning, the tissue was infiltrated with 30% sucrose solution, frozen, and sectioned in a cryostat (Leica, Wetzlar, Germany) at 40 μm. The prelimbic cortex was sectioned in the sagittal plane, and the LEC was sectioned in the coronal plane. Sectioned tissue was stained with cresyl violet and imaged under a light microscope to locate electrode and infusion cannulae locations. All tetrodes containing neurons were located in the prelimbic region and were included in the analyses.

## RESULTS

### Behavioral performance

During each daily session, four male Long-Evans rats underwent multiple epochs of trace eyeblink conditioning in the same chamber (Figure 1A). The initial acquisition phase included two epochs (Figure 1B), in which the rats first received 20 presentations of the conditioned stimulus (CS; tone or light, 100 msec) by itself (CS-alone block; Figure 1A), followed by 50 pairings of the CS and mildly aversive eyelid shock (US) over a 500-msec interval (CS-US block). This block design enables rats to adjust their behavioral responses to the same CS according to the temporal context governing the CS-US contingency (Morrissey et al., 2017; Pilkiw et al., 2017). Also, the CS-alone block serves as a control to test how task demands affect the stability of neural firing patterns. The CS presentations were separated by inter-trial intervals ranging from 20 to 40 seconds. Across 5-7 daily sessions, all rats developed anticipatory blinking responses that peaked near the expected onset of the US (conditioned response, CRs). In parallel, they also learned to withhold their eyeblink responses in the CS-alone blocks.

**Figure 1.**
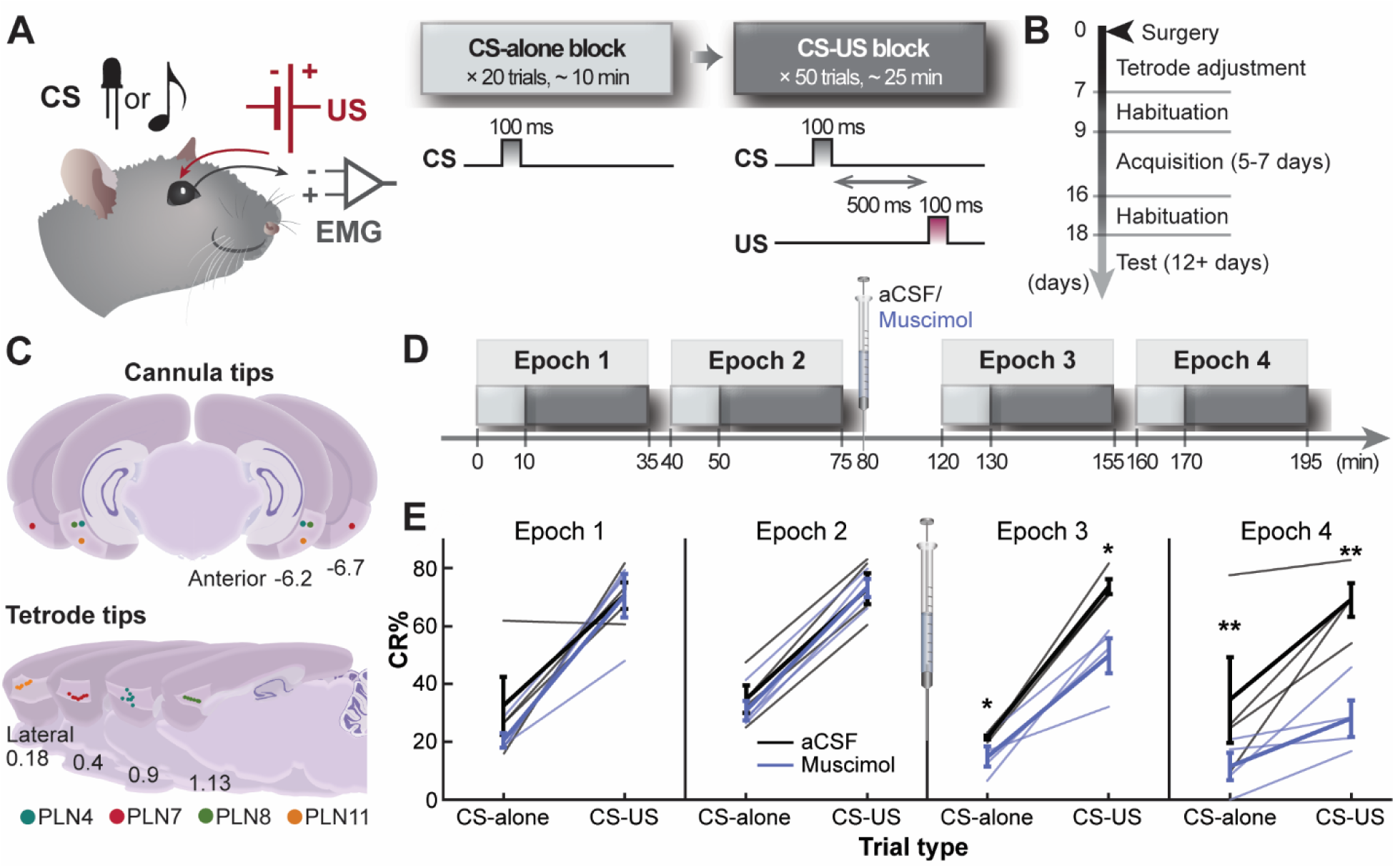
Task schedule and the effect of LEC inhibition on behavior. **A.** The sequence of stimulus presentations in each epoch. **B.** The experimental schedule. **C.** The schematic representations of cannula location and the site of single-unit recording. **D.** The timeline of each test session including four identical conditioning epochs (Epoch 1-4) and the pharmacological manipulation. **E.** The proportion of trials in which rats expressed conditioned responses (CR%; n = 4 rats, mean ± s.e.m.). **\*** and ** indicate p < 0.05 and 0.01 in two-way repeated-measures ANOVA followed by *post hoc* Tukey HSD.

Once the rat expressed CRs in 60% of CS presentations in at least one of the CS-US blocks, the rats moved onto the test phase in which the lateral entorhinal cortex (LEC) was inactivated by bilateral infusions of the GABA_A_ receptor agonist, muscimol (Figure1B, C). We chose this approach over optogenetic and chemogenetic approaches to attain robust impairments in memory retrieval (Morrissey et al., 2012). Each test session consisted of four iterations (Epoch 1-4) of the same sequence of the CS-alone and CS-US blocks in the same conditioning chamber (Figure 1D). Either the original tone or light stimulus was used as the CS throughout the four epochs in each session. After Epoch 2, artificial cerebral spinal fluid (aCSF; 7 sessions across four rats) or muscimol (11 sessions) was bilaterally infused through chronically implanted microinfusion cannulae in the LEC. All rats underwent at least one session with each injection type in a pseudo-randomized order (Table 1). The post-mortem histological analysis confirmed that all cannulae were located in the dorsolateral part of the LEC (Figure 1C), from which monosynaptic projections to the prelimbic region (PrL) originate (Kerr et al., 2007). Based on our previous work that estimated the spread of muscimol conjugated with fluorescent molecules (Morrissey et al., 2012), the volume of drug solution was set to inhibit neurons in both superficial and deep layers of the LEC.

Consistent with our previous work (Morrissey et al., 2012), pharamacological inactivation of the LEC decreased the proportion of trials with CRs (CR%) in the CS-alone and CS-US blocks (Figure 1E; three-way repeated measures ANOVA, n = 4 rats, Drug×CS type×Epoch interaction, F_3, 9_ = 1.552, p = 0.268; Drug×Epoch interaction, F_3, 9_ = 5.525, p = 0.020; CS type×Epoch, F_3, 9_ = 1.958, p = 0.191; Drug×CS type, F_3, 9_ = 1.437, p = 0.317). After collapsing CS type, we found that in the two epochs after the injection, CR% was lower in muscimol than aCSF sessions (follow-up paired t-test; Epoch 3, p = 0.015; Epoch 4, p = 0.007). This was not the case in the epochs before the injection (Epoch 1, p = 0.351; Epoch 2, p = 0.573). In addition, in muscimol, but not aCSF, sessions, CR% changed significantly across the four epochs (follow-up one-way repeated measures ANOVA, Muscimol, F_3, 21_ = 11.503, p < 0.001; aCSF, F_3, 21_ = 0.277, p = 0.841). CR% after the infusion was lower than that before the infusion (follow-up paired t-test vs. Epoch 2; Epoch 3, p = 0.005; Epoch 4 p = 0.001) though CR% did not differ between the two epochs after the infusion (p = 0.101). In parallel, even in muscimol sessions, CR% was higher in the CS-US block than the CS alone block (a main effect of CS type, F_1, 3_ = 261.381, p < 0.001), suggesting that LEC inhibition impaired the retrieval of the CS-US association rather than paralizying eyelid muscles.

### High task demands suppress drift in PrL ensemble firings

During the test sessions, we extracellularly recorded action potentials of neurons in the PrL with a chronically implanted mechanical drive (Kloosterman et al., 2009) containing 14 independently movable, four-channel electrodes (tetrodes; Wilson and McNaughton, 1993). The final locations of the tetrodes were evenly distributed along the anterior-posterior axis within the PrL (Figure 1C). We first investigated to what degree PrL ensembles exhibited consistent activity patterns across repeated experiences and how the consistency changed depending on task demands. To this end, we analyzed the activity of all 1067 neurons recorded in all 18 sessions (regardless of drug assignment) during the two epochs before the infusion (*i.e.*, Epochs 1 and 2; Figure 1D).

Consistent with our previous findings (Takehara-Nishiuchi and McNaughton, 2008; Morrissey et al., 2017), a large proportion of PrL neurons changed their firing rates in response to the CS and US presentation (Figure 2A, Bin size = 100 msec, Cells × 16 Bins, CS onset at the 4^th^ bin). In the CS-US block, most of these stimulus-responding neurons in Epoch 1 showed consistent firing patterns in Epoch 2 (Figure 2Aa). In contrast, ensemble patterns in the CS-alone block were less similar between the two epochs (Figure 2Ab). The Pearson correlation coefficient of these firing rate matrices was higher in the CS-US block than the CS-alone block (Figure 2B; n = 20 runs with 300 subsampled cells, paired t-test, p < 0.001).

**Figure 2.**
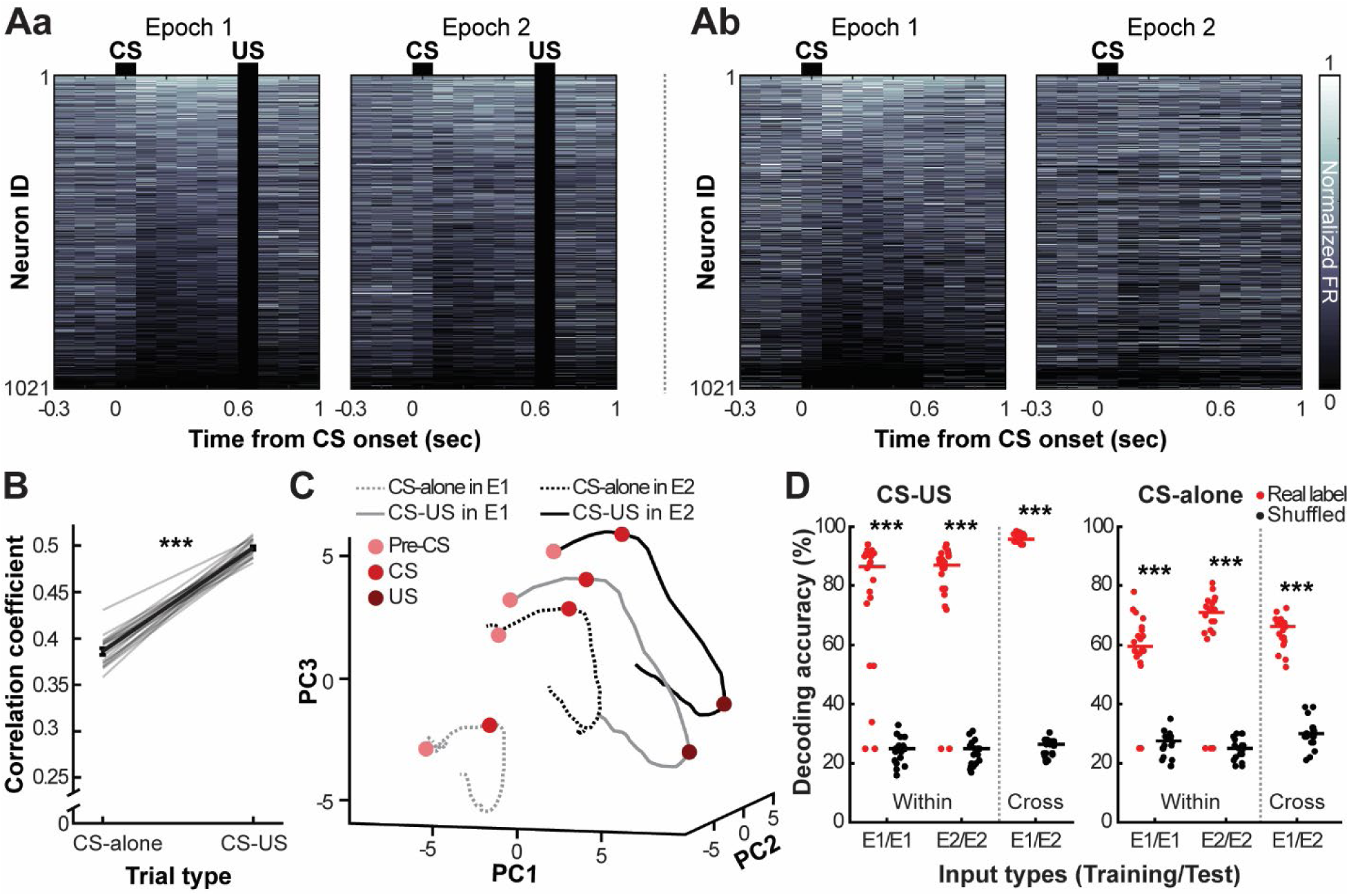
Stabilization of PrL ensemble firings by high task demands. **A.** Greyscale plots showing the normalized firing rate (max = 1 in each neuron) during two epochs before the drug infusion (a, CS-US block; b, CS-alone block). Neurons were sorted based on the CS-induced firing rate during the first epoch, from the largest increase (Neuron #1) to the largest decrease (Neuron #1005). Black bars on the top indicate the CS, while black rectangles mask US artifacts. **B.** The Pearson correlation coefficient of ensemble firing rates between two epochs (n = 20 runs with 300 subsampled neurons, mean ± s.e.m.). **C.** Trajectories of ensemble firing patterns projected onto the top three principal components (PC). **D.** Accuracy of decoding of the task events from ensemble firing rates in trials in the same epoch (Within) or different epochs (Cross). Each dot represents decoding accuracy in each run with 300 subsampled neurons. Horizontal bars indicate the median. ***, p < 0.001 in Wilcoxon signed-rank tests.

To better visualize the change in neural representations across the epochs, we then examined the time-evolving trajectory of ensemble activity in neural state space (Raposo et al., 2014; Remington et al., 2018; Russo et al., 2020). Principal component analysis (PCA) was applied to ensemble firing rate matrices during a 2.6-sec window centering on CS onset (excluding the 100-msec US period) to project the time-varying ensemble firing patterns in neural state space (Figure 2C). In the CS-US block, ensemble activity formed a smooth trajectory that exited the baseline state (Pre-CS, light red circle) upon CS presentations (CS, red circle), and entered an orbit smoothly transitioning until the US presentation (US, dark red circle). The geometry of the trajectory was consistent in Epochs 1 (solid grey line) and 2 (solid black line). In the first CS-alone block, the CS presentation induced a similar change in the trajectory, followed by a smooth transition that resembled the trajectory in the CS-US block, albeit a smaller orbit (grey dotted line). The orbit appeared to be enlarged in Epoch 2 (black dotted line) and displaced to a region closer to the trajectories during the CS-US blocks.

To quantify the difference in these neuronal trajectories and their consistency across the two epochs, we applied support vector machine classifiers to ensemble firing rates during four task phases separately in CS-alone and CS-US blocks. These phases were the pre-CS (-300-0 ms from CS onset), CS (0-300 ms), CS-US interval (300-600 ms), and post-US (0 -300 ms from US offset) periods. We included CS-US interval and post-US phases in the CS-alone block analyses because the neural trajectory in the CS-alone block followed similar trajectories to that in the CS-US block (Figure 2C). Classifiers were first trained with firing rates from a subset of trials and then tested them on the remaining trials in each epoch (within-epoch decoding). Better performance of the classifier reflects more accurate encoding of the task events. In the CS-US block, classifiers decoded the four task phases better than chance (Figure 2D, within; n = 20 runs with 300 subsampled neurons; Wilcoxon signed-rank test vs. shuffled labels, in both epochs, ps < 0.001). Similarly, in the CS-alone block, the classifier successfully discriminated the task phases from one another better than chance (in Epochs 1 and 2, ps < 0.001). To test the consistency of this ensemble activity across the two epochs, we then trained classifiers with all the trials in Epoch 1 and tested on all the trials in Epoch 2 (cross-epoch decoding). Consistent with the high correlation between the two ensemble firing rate matrices in the CS-US block (Figure 2B), decoding accuracy was better than chance (Figure 2D, cross; p < 0.001). This observation suggests that neuronal ensembles stably maintained information about the task events across the two epochs. Cross-epoch decoding accuracy was also better than chance in the CS-alone block (p < 0.001), suggesting that despite the low correlation of fine-scale ensemble firing patterns (Figure 2B) and the major displacement of the neuronal trajectory (Figure 2C), neuronal ensembles held the CS information with sufficient stability across the epochs.

As an alternative approach for quantifying the similarity of ensemble firing patterns, we also calculated Pearson correlation coefficients between the two firing rate matrices (300 cells × 4 task phases) constructed from trials within each epoch or between two epochs. In CS-US blocks, correlation coefficients with real data (Epoch 1, 0.555 ± 0.006; Epoch 2, 0.522 ± 0.007; Cross, 0.632 ± 0.006) were greater than those with shuffled data (Epoch 1, -0.005 ± 0.007; Epoch 2, 0.003 ± 0.006; Cross, -0.009 ± 0.005; paired t-test, all ps < 0.001). Compared to the CS-US block, correlation values were lower in the CS-alone block (Epoch 1, 0.370 ± 0.007; Epoch 2, 0.378 ± 0.006; Cross, 0.378 ± 0.005; all ps < 0.001); however, they were still significantly higher than those with shuffled data (Epoch 1, -0.004 ± 0.005; Epoch 2, 0.006 ± 0.007; Cross, 0.001 ± 0.006; all ps < 0.001).

Collectively, these results suggest that PrL neuronal ensembles maintained their consistent selectivity for the stimulus sequences across repetitions; however, the fine-scale ensemble firing patterns were more stable for the behaviorally relevant sequences than the incidental sequences.

### LEC inhibition destabilizes PrL ensemble representations of learned stimulus associations

We then examined how LEC inhibition affected the stability of PrL ensemble representations by applying the same analytical approaches to the ensemble activity before and after the infusion (*i.e.*, Epoch 2 vs. Epoch 3). After aCSF infusions, the PrL maintained consistent ensemble activity in the CS-US block (Figure 3Aa), though the ensemble patterns appeared less stable in the CS-alone block (Figure 3Ab). In contrast, muscimol infusions induced a drastic change in ensemble patterns in both the CS-US (Figure 3Ac) and CS-alone blocks (Figure 3Ad). In both blocks, the correlation coefficient between the two firing matrices was lower in muscimol than in aCSF sessions (Figure 3B; two-way mixed ANOVA, Block×Drug, F_1, 38_ = 0.054, p = 0.818; Block, F_1, 38_ = 1362.223, p < 0.001; Drug, F_1, 38_ = 107.856, p < 0.001).

**Figure 3.**
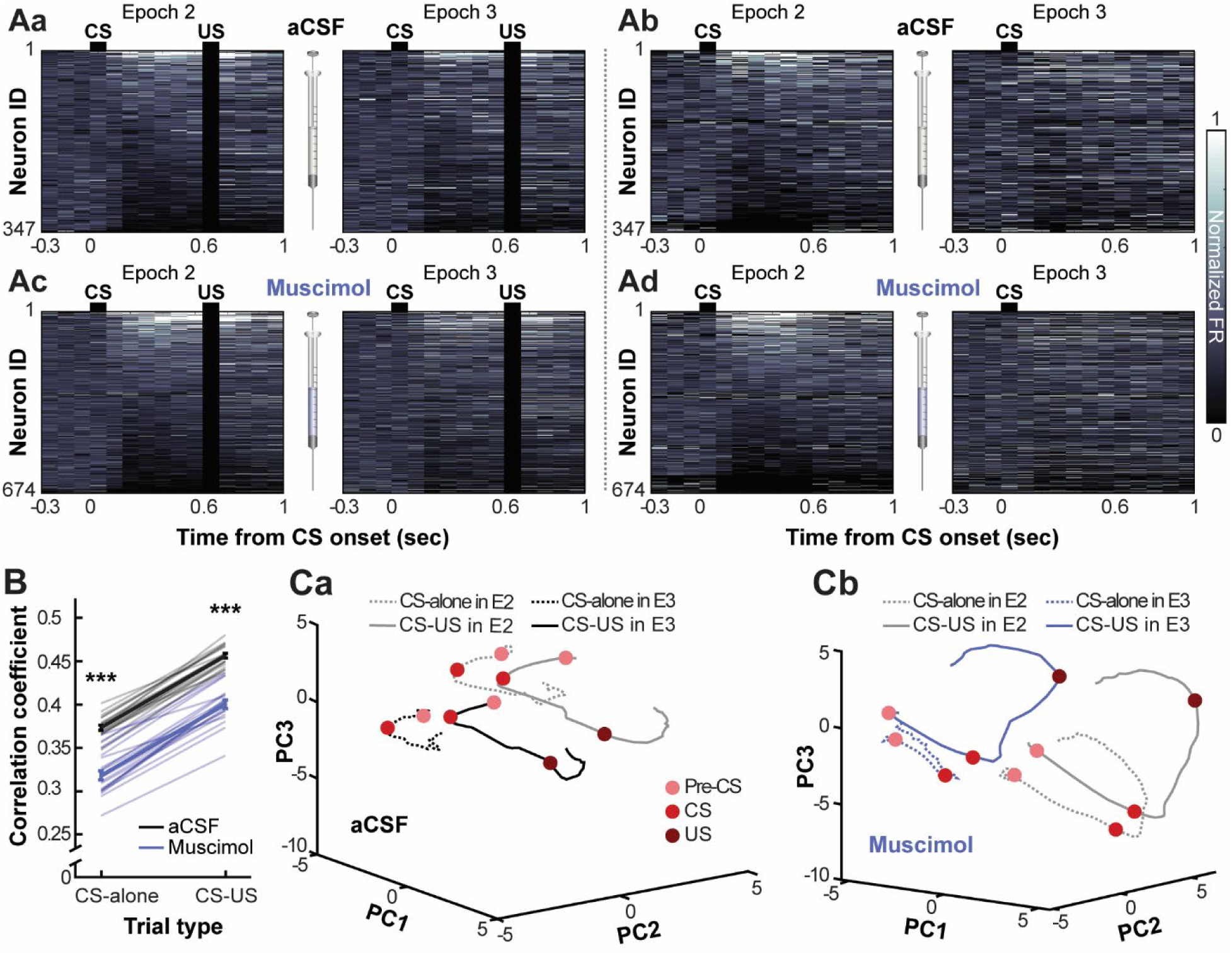
Reconfiguration of PrL ensemble representations of task events following LEC inhibition. **A.** Greyscale plots showing the normalized firing rate (max = 1 in each neuron) during the epochs before and after aCSF (a, b) or muscimol infusions (c, d). Neurons were sorted based on the CS-induced firing rates during the epoch before the infusion. Black bars on the top indicate the CS, while black rectangles mask US artifacts. **B.** The correlation coefficient of ensemble firing rates between two epochs before and after the infusion. (n = 20 runs with 300 sub-sampled cells, mean ± s.e.m). ***, p < 0.001 in two-way mixed ANOVA followed by t-tests. **C.** Trajectories of ensemble firing patterns projected onto the top three principal components (PC) in sessions with aCSF (a) and muscimol infusions (b).

In neural state space, aCSF infusions only marginally affected the location and geometry of the neural trajectory (Figure 3Ca), whereas muscimol infusions moved the trajectory farther away from the original location (Figure 3Cb). Despite this major displacement, the geometry of the neural trajectory in the CS-US block was mostly unaffected. This observation suggests that the PrL generated a new ensemble firing pattern that accurately encoded the four task phases. In the CS-alone block, after the CS, the neural trajectory immediately returned to the pre-CS state instead of following a smooth orbit like that in the CS-US block.

Consistent with these visual impressions, the accuracy of the within-epoch decoding in the CS-US block was comparable between muscimol and aCSF sessions (Figure 4Aa; within; n = 20 runs with 300 subsampled cells; Wilcoxon rank-sum test, Epoch 2, p = 0.455; Epoch 3, p = 0.626). In contrast, the accuracy of the cross-epoch decoding was lower in muscimol than in aCSF sessions (p < 0.001). Examination of the specific classification errors (the ‘confusion matrix,’ Figure 4Ba) showed that most of the inaccurate classifications were due to errors in discriminating the CS-US interval period (included in the “CS” and “CS-US interval” phases) from the other task phases. To confirm this observation statistically, we quantified decoding accuracy separately for firing patterns taken from each of the four task phases. Muscimol sessions showed lower decoding accuracy for the CS and interval period than aCSF sessions (Figure 4Ca, Wilcoxon rank-sum test, ps < 0.001). In contrast, decoding accuracy for the pre-CS period was higher in muscimol than in aCSF sessions (p < 0.001). Muscimol infusions did not affect decoding accuracy for the post-US period (p = 0.089). In the CS-alone block, the within-epoch decoding accuracy was lower in muscimol than in aCSF sessions after, but not before, the infusion (Figure 4Ab; Epoch 2, p = 0.232; Epoch 3, p = 0.001). Classifiers frequently decoded two task phases after CS onset as the pre-CS period (Figure 4Bb), echoing the collapsed neural trajectory following muscimol infusions (Figure 3Cb). Furthermore, cross-epoch decoding accuracy was also lower in muscimol than in aCSF sessions (p < 0.001; Figure 4Bb). Muscimol infusions lowered the accuracy of decoding firing patterns during the CS (p = 0.002) as well as the periods matching the interval (p < 0.001) and post-US period (p = 0.005) of the CS-US block. But, they did not affect decoding accuracy for the pre-CS period (p = 0.324; Figure 4Cb).

**Figure 4.**
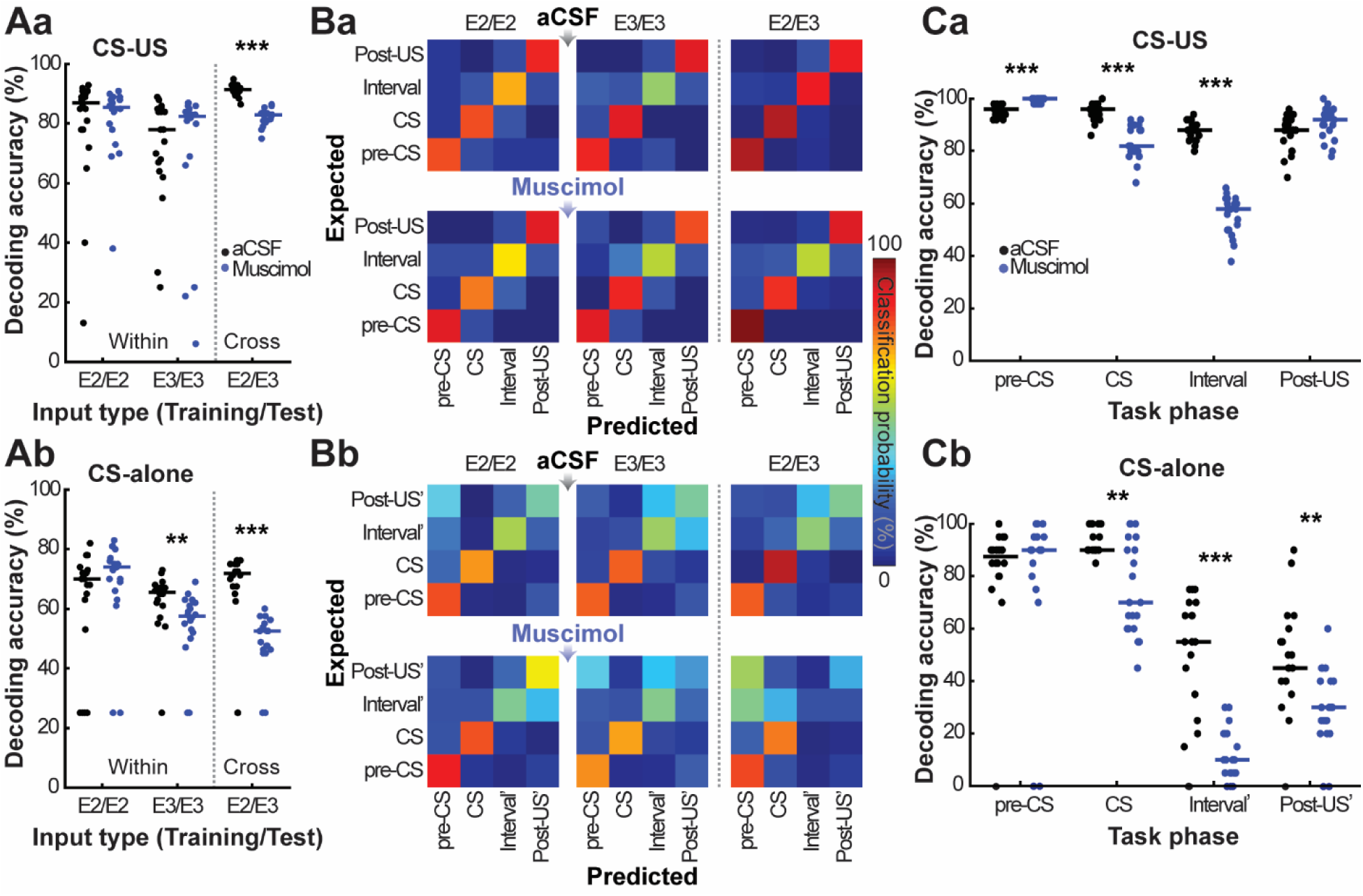
Impaired reinstatement of ensemble firing patterns by LEC inhibition. **A.** Decoding accuracy of the task events within each epoch (Within) or between two epochs (Cross) in the CS-US block (a) and the CS-alone block (b). Each dot represents decoding accuracy in each of 20 runs with 300 subsampled neurons. Horizontal bars indicate the median. **, ***, p < 0.01, 0.001 in Wilcoxon rank-sum tests. **B.** Confusion matrices indicating proportions of task events that were identified by the classifier correctly (warm colors along the diagonal) or misidentified as a different event (lighter colors off-diagonal) in the CS-US block (a) and the CS-alone block (b). The expected task phase (y-axis) shows the task phase from which a firing rate pattern was extracted. The predicted task phase (x-axis) shows which task phase the decoder classified the firing rate pattern as. The color indicates the probability of firing patterns in each task phase being categorized in one of the four task phases. Although the US was not delivered during the CS-alone block, firing rates were calculated during time windows corresponding to the interval (Interval’) and post-US period (Post-US’) in the CS-US block. **C.** Decoding accuracy in cross-epoch decoding analyses calculated separately for each of the four task phases in the CS-US block (a) and the CS-alone block (b), as in A.

Collectively, these findings suggest that LEC inhibition marginally affected PrL ensemble selectivity for the task phases within each epoch; however, it destabilized the ensemble activity across the epochs mainly by preventing the CS inputs from reinstating the original network state that bridges the temporal gap between the CS and US.

### LEC inhibition induces various changes in stimulus-evoked firings of individual PrL neurons without affecting the population-level stimulus representations

We next examined the impact of LEC inhibition on the firing patterns of individual neurons (Figure 5). After muscimol infusions, some neurons became completely silent (Neuron 1) or maintained their baseline firings but lost stimulus-evoked responses (Neuron 2; FR loss). In contrast, others increased their baseline firing rates and acquired firing responses to the CS (Neurons 3, 4; FR gain). Similarly, some neurons lost their differential firing responses between the CS presented alone and the CS paired with the US (Neurons 5, 6; Selectivity loss), while others gained the differential responses (Neurons 7, 8; Selectivity gain). Firing patterns of other neurons barely changed after the infusion (Neurons, 9, 10; Stable).

**Figure 5.**
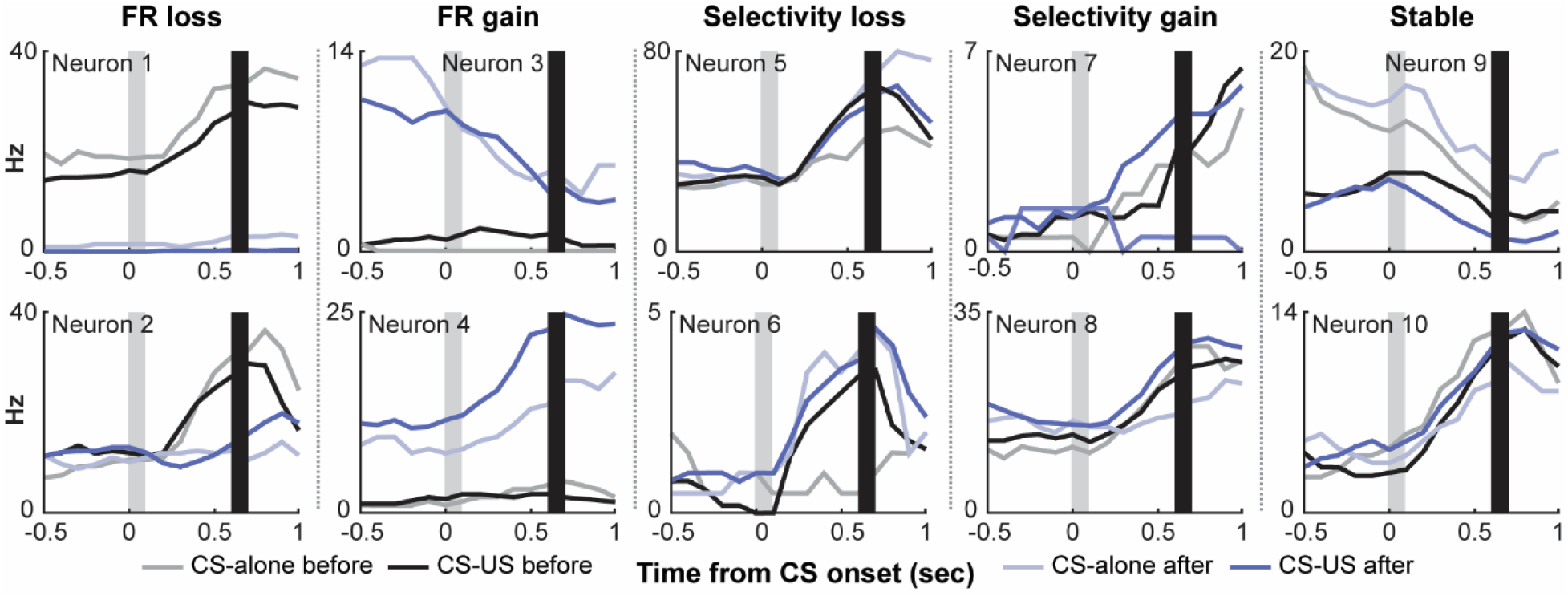
Diverse effects of LEC inhibition on firing patterns of individual PrL neurons. Examples of firing patterns of individual neurons during the epochs before (black) and after (blue) muscimol infusions. The lines in light and dark color show the firing pattern in the CS- alone and CS-US block, respectively. Light grey bars indicate the CS, and black bars mask US artifact.

In the CS-US block, ∼5 and ∼10% of neurons increased and decreased their baseline firing rates after aCSF infusions, respectively (Figure 6A; p < 0.05 in random permutation tests). Muscimol infusions increased the proportion of these neurons with unstable baseline firing rates (Figure 6A, ∼25%; binominal test, p < 0.05). Due to the bidirectional change, however, the effect of muscimol infusions on the overall distribution of baseline firing rates was marginal with the slight increase in neurons with decreased baseline firing rate (Figure 6B, insets; Wilcoxon rank-sum test, p = 0.023).

**Figure 6.**
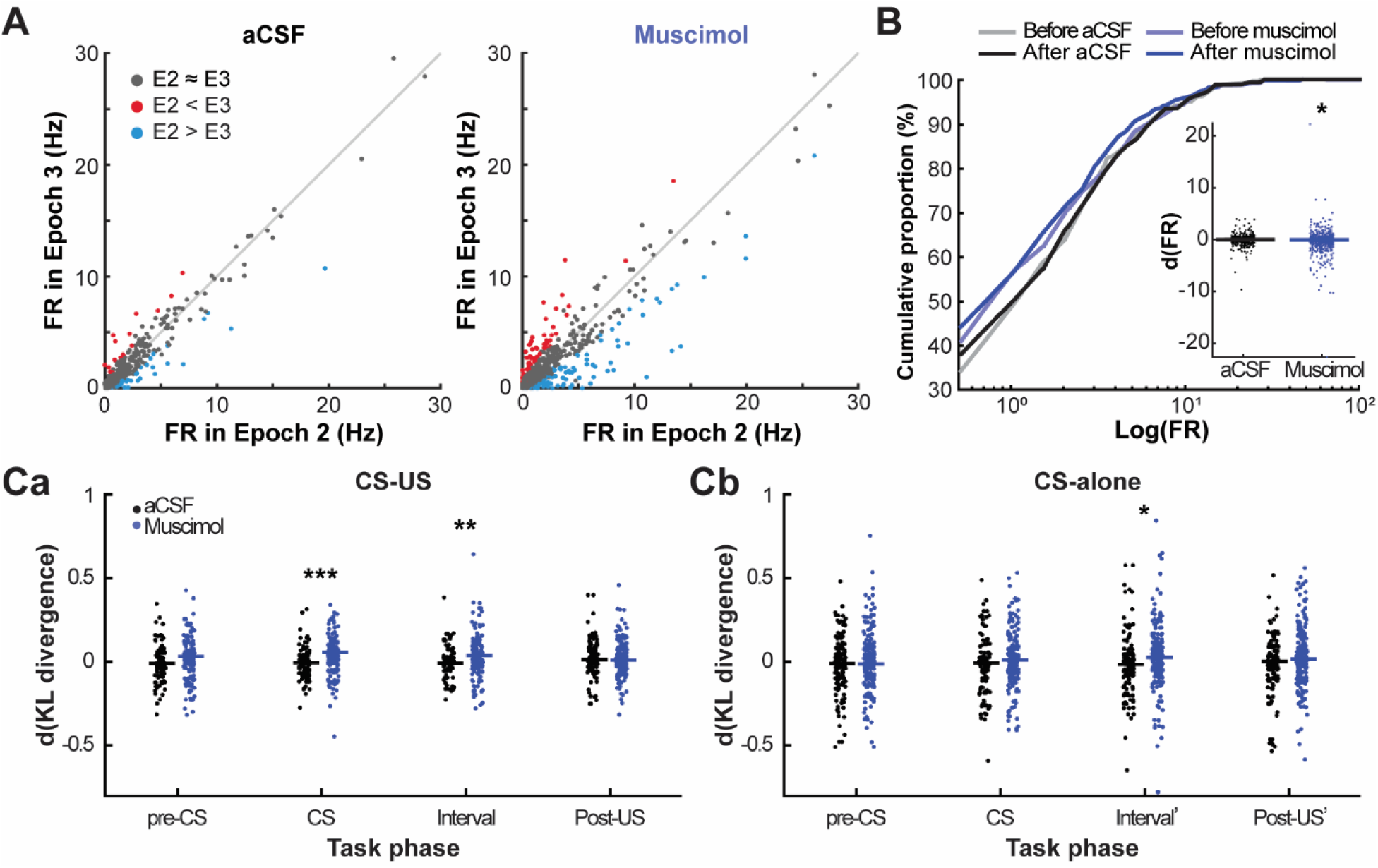
Effects of LEC inhibition on spontaneous firing rates and across-trial firing stability of PrL individual neurons. **A.** Spontaneous firing rate (FR) of each neuron before (x-axis) and after the infusion (y-axis; one dot per neuron, color indicates the result of random permutation tests, p < 0.05) in aCSF and muscimol sessions. **B.** The cumulative distributions of spontaneous FR. The inset shows the difference in spontaneous FRs between before and after the infusion (one dot per neuron). **C.** The difference in a measure quantifying the stability of firing rates across trials before and after the infusion (one dot per neuron). Firing rates were calculated during each of the four task phases. The horizontal bars indicate the median. *, **, *** indicate p < 0.05, 0.01, 0.001 in Wilcoxon rank-sum test.

We then examined whether LEC inhibition affected the stability of firing rates within each epoch by quantifying the degree to which the distribution of firing rates across trials diverged from a uniform distribution (see Materials and methods). The greater divergence indicates a more significant drift of firing rates within each epoch. In the CS-US block, muscimol infusions worsened the across-trial drift of firing rates during the CS and intervals (Figure 6Ca, Wilcoxon rank-sum test, p < 0.001, 0.007), but not during the pre-CS or post-US period (p = 0.066, 0.798). In the CS-alone block, muscimol infusions only exaggerated the across-trial drift of firing rates during the period matching the interval of the CS-US block (p = 0.027; other phases, p > 0.05). These results suggest that the integrity of LEC is necessary for neurons to show consistent firing responses to repeated presentation of the CS particularly when it was predictive of the US.

Next, we investigated the effect of LEC inhibition on the proportion of neurons responding to the stimuli. In both aCSF and muscimol sessions, ∼30% of neurons responded to the CS, and the proportion of these CS-responding neurons was comparable before (Epoch 2) and after (Epoch 3) the infusion (Figure 7Aa). After aCSF infusion, ∼60% of CS-coding neurons maintained the same response type (*i.e.*, firing rate increase or decrease; Figure 7Aa, grey arrows; Figure 7Ab, Retention), while ∼20% of the non-responding neurons gained a response (orange arrows). The latter, newly recruited neurons consisted of ∼45% CS-responding neurons after the infusion (Figure 7Ab, Recruitment). These proportions were not affected by muscimol infusions (binominal test, p > 0.05). In addition, muscimol infusions did not affect the overall distribution of the magnitude of CS-evoked firing responses (Figure 7Ac; the change after the infusion, Wilcoxon rank-sum test, p = 0.500).

Compared to the CS-evoked responses, the US-evoked responses appeared more stable across the epochs: for neurons that responded to the US before the infusion, ∼70% maintained their US-evoked responses after the infusion (Figure 7Ba), while ∼20% of the non-responding neurons gained a response. Muscimol infusions decreased the retention rate but increased the reinstatement rate (Figure 7Bb, binominal tests, ps < 0.05). As a result, the proportion of the US-responding neurons and the magnitude of responses were comparable before and after the infusion (Figure 7Bc; the change after the infusion, Wilcoxon rank-sum test, p = 0.484).

**Figure 7.**
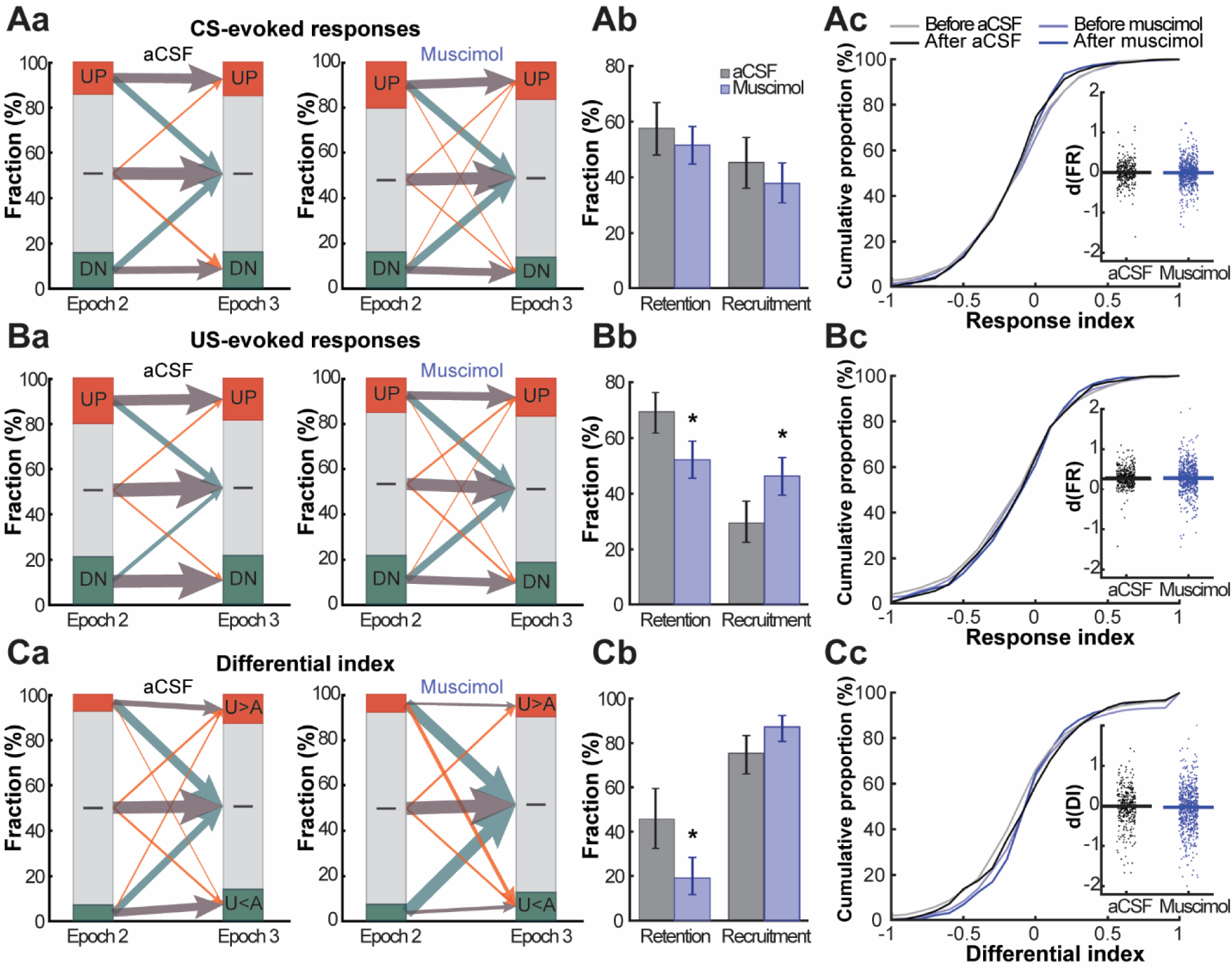
Effects of LEC inhibition on the sparsity and consistency of task event representations in the PrL. **A.** (a) In aCSF and muscimol sessions, neurons were categorized into those with a significant FR increase (UP), decrease (DN), or no change (—) in response to the CS (p < 0.05 in random permutation tests). The width of arrows indicates the fraction of neurons that changed their category after the infusion while their color indicates the type of change (grey, no change; green, loss of significance; orange gain of significance). (b) For neurons with significant CS-evoked FR responses before the infusion, the fraction of neurons maintained the same type of responses after the infusion (Retention). For neurons with significant CS-evoked FR responses after the infusion, the fraction of neurons that gained the responses after the infusion (Recruitment). Error bars indicate 95% confidence intervals. (c) The cumulative distribution of the magnitude of CS-evoked firing responses in the CS-US block before and after the infusion. The magnitude was quantified by the response index (RI) that becomes 0 for no change and -1/1 for the most robust decreased/increased firing rates. The inset shows the change in the response magnitude after the infusion (one dot per cell). The horizontal bars indicate the median. **B.** The same for US-evoked FR responses. **C.** (a) In aCSF and muscimol sessions, neurons were categorized into those with significantly higher CS-evoked FR rates in the CS-US than CS-alone block (U>A), significantly lower rates (U<A), or no difference (—; p < 0.05 in random permutation tests). The definition of the arrow width and color is the same as in A. (b) For neurons with significant FR differentiation before the infusion, the fraction of neurons that maintained the same FR differentiation after the infusion (Retention). For neurons with significant FR differentiation after the infusion, the fraction of neurons that gained the differentiation after the infusion (Recruitment). Error bars indicate 95% confidence intervals. (c) The cumulative distribution of the degree of differentiation of CS-evoked firing responses between the CS-US block and CS-alone block. The differentiation was quantified by the differential index (DI) that becomes 0 for no differentiation and -1/1 for lower/higher firing rates in the CS-US block than CS-alone block. The inset shows the change in the firing differentiation from before to after the infusion (one dot per cell). The horizontal bars indicate the median.

Consistent with our previous works (Takehara-Nishiuchi and McNaughton, 2008; Morrissey et al., 2017; Takehara-Nishiuchi et al., 2020), only a subset of CS-responding neurons differentiated CS-evoked firing rates between the CS-alone and CS-US block (Figure 7Ca). After aCSF infusion, ∼45% of these association-selective neurons maintained their selectivity (Figure 7Cb). Notably, ∼20% of the non-selective neurons gained selectivity after the infusion, leading to an apparent increase in the proportion of associating-selective neurons. After muscimol infusions, most association-selective neurons lost their selectivity (Figure 7Ca), and their retention rate was significantly lower than after aCSF infusions (Figure 7Cb; binominal test, p < 0.05). In parallel, ∼20% of the non-selective neurons gained selectivity, while 20% of the neurons reversed their directionality of firing differentiation (*i.e.*, higher or lower firing rates in the CS-US block than in the CS-alone block). As such, the recruitment rate showed a trend toward an increase (Figure 7Cb). Despite these changes, the overall distribution of the selectivity for the CS-US association was not affected by either infusion type (Figure 7Cc; the change after the infusion, Wilcoxon rank-sum test, p = 0.118).

In the CS-alone block, muscimol infusions had negligible effects on these measures: there were no significant differences in the infusion-induced change in baseline firing rates (data not shown; Wilcoxon rank-sum test, p = 0.823) or CS-evoked firing responses (data not shown; p = 0.543) between muscimol and aCSF sessions.

Collectively, these findings suggest that although the firing patterns of individual neurons were markedly reconfigured across repetitions, the overall proportion of neurons exhibiting the stimulus-evoked responses or the selectivity of their firings for stimulus associations were stably maintained. The LEC inhibition impaired the reactivation of these stimulus-responding and association-selective neurons but did not prevent other neurons from gaining stimulus selectivity. As such, the LEC inhibition did not have major effects on the selectivity for learned stimulus associations at the population level.

### LEC inhibition abolishes the beneficial effect of task demands on PrL ensemble stability

Our behavioral task included two epochs after the infusions (*i.e*., Epochs 3 and 4), which enabled us to compare ensemble stability over time with and without the integrity of the LEC. As in the previous comparisons (Figure 2A, 3A), in aCSF sessions, comparable ensemble patterns were detected in Epochs 3 and 4 with greater stability in the CS-US block than in the CS-alone block (Figure 8Aa, b). In contrast, the ensemble patterns in muscimol sessions appeared dissimilar between Epochs 3 and 4 in both blocks (Figure 8Ac, d). Notably, muscimol infusions lowered the correlation coefficient of ensemble firing matrices in the CS-US block (Figure 8B; n = 20 set of 300 sub-sampled cells; two-way mixed ANOVA, Block×Drug, F_1, 38_ = 166.223, p < 0.001; follow-up t-test, p < 0.001) but not in the CS-alone block (p = 0.255), suggesting that LEC inhibition selectively eliminated the beneficial effects of high task demands on the stability of fine-scale ensemble firings.

**Figure 8.**
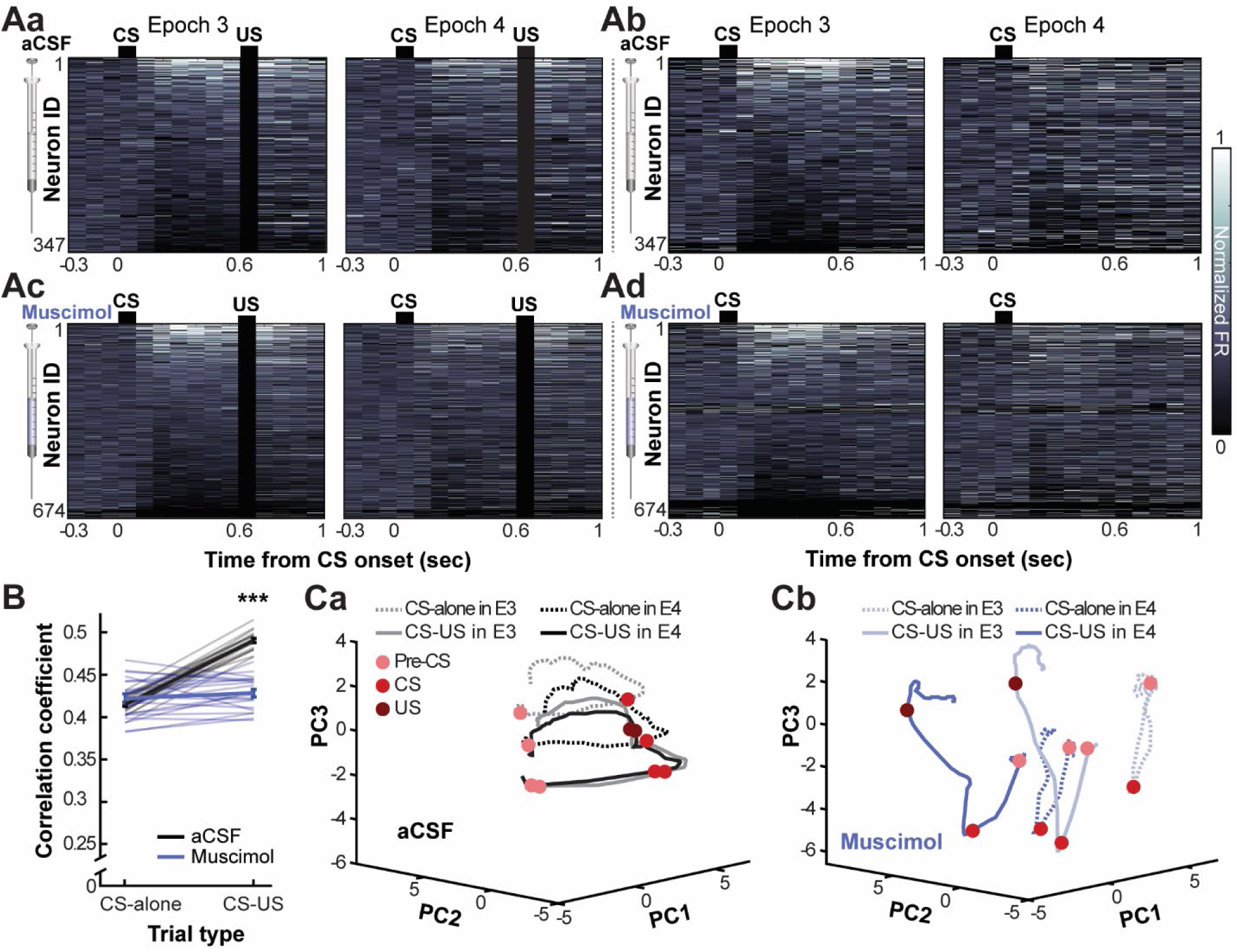
Loss of beneficial effects of high task-demand on PrL stability after LEC inhibition. **A.** Greyscale plots show the normalized firing rate during the two epochs after aCSF (a, b) or muscimol infusions (c, d). Neurons were sorted based on the CS-evoked firing rate during the preceding epoch. Black bars on the top indicate the CS, while black rectangles mask US artifacts. **B.** The correlation coefficient of ensemble firing rates between two epochs after the infusion (n = 20 runs with 300 sub-sampled cells, mean ± s.e.m). ***, p < 0.001 in two-way mixed ANOVA followed by t-test. **C.** Trajectories of ensemble firing patterns projected onto the top three principal components (PC).

In parallel, the neuronal trajectory in muscimol sessions became shifted to a new location in neural state space as the trial blocks and epochs progressed (Figure 8Cb). Such representational drift appeared smaller in aCSF sessions (Figure 8Ca). In both infusion types, the geometry of the neural trajectory was similar between the two epochs.

Consistent with these observations, ensemble decoding analyses revealed that muscimol infusions did not affect the within-epoch decoding accuracy in the CS-US block of Epoch 4 (Figure 9Aa; Wilcoxon rank-sum test, p = 0.569). However, the accuracy of the cross-epoch decoding was significantly lower in muscimol than in aCSF sessions (p < 0.001). Most of the inaccurate classifications were due to errors in discriminating the CS-US interval from the pre-CS periods (Figure 9Ba). Muscimol infusions lowered decoding accuracy for the CS (Figure 9Ca, p < 0.001) and interval periods (p < 0.001), while they improved the decoding accuracy for the pre-CS (p < 0.001) and post-US periods (p = 0.009). In the CS-alone block, muscimol infusions lowered the accuracy of the within- (Figure 9Ab, p < 0.001) and cross-epoch decoding (p < 0.001). Inaccurate cross-epoch decoding originated from errors in the decoding of the two task phases after CS offset (the period corresponding to the interval and post-US period of the CS-US block; Figure 9Bb, Cb). Muscimol infusions lowered the accuracy of decoding for the pre-CS (p = 0.005) and the two subsequent periods (both ps < 0.001), but not the CS period (p = 0.158).

**Figure 9.**
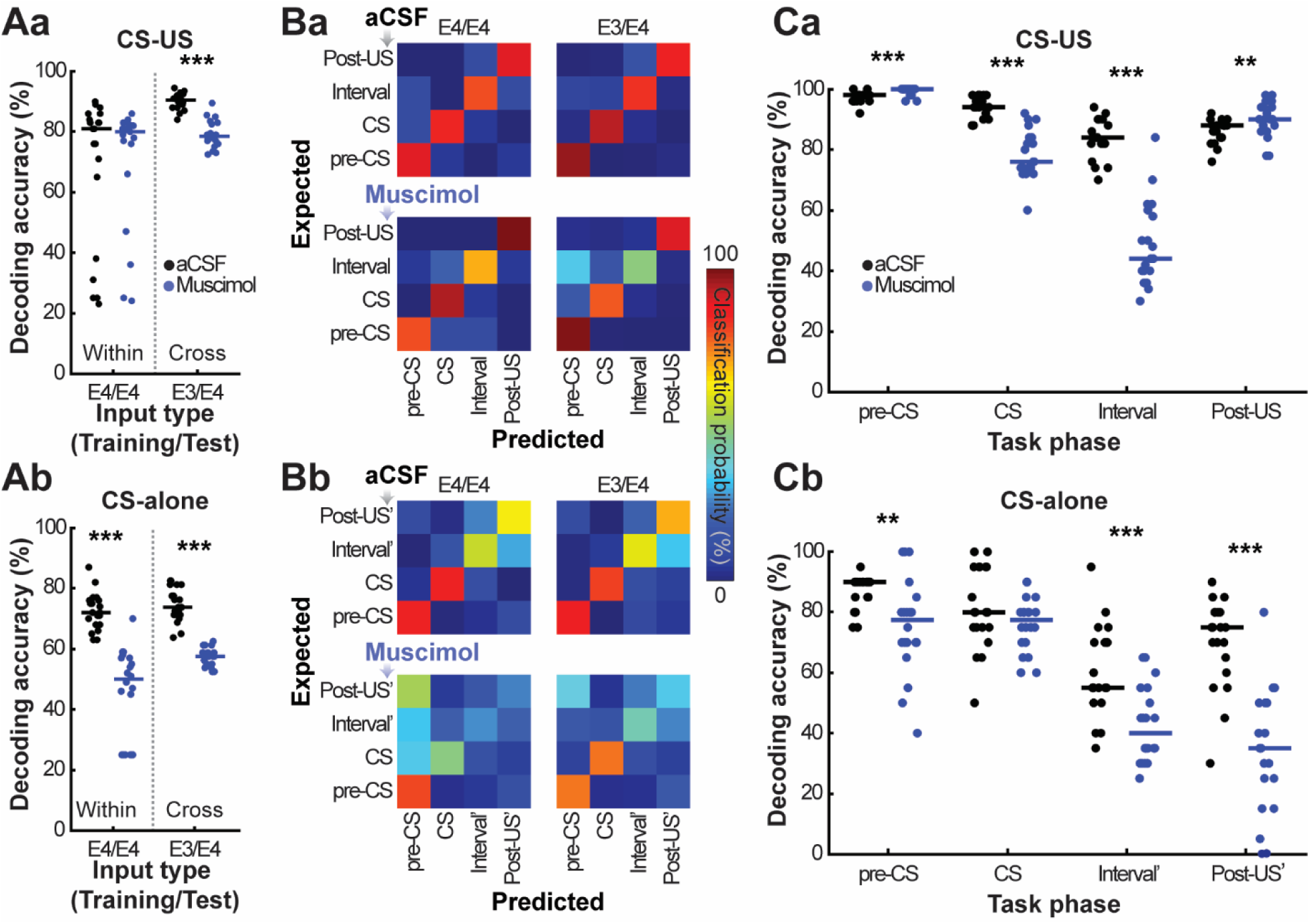
Exaggerated drift of PrL ensemble firing patterns by LEC inhibition. **A.** Decoding accuracy of the task events from ensemble firing rates in trials in the same epoch (Within) or different epochs (Cross) in the CS-US block (a) and the CS-alone block (b). Each dot represents decoding accuracy in each of 20 runs with 300 subsampled cells. Horizontal bars indicate the median. *, **, ***, p < 0.05, 0.01, 0.001 in Wilcoxon rank-sum tests. **B.** Confusion matrices indicating proportions of task events that were identified by the classifier correctly (warm colors along the diagonal) or misidentified as a different event (lighter colors off-diagonal) in the CS-US block (a) and the CS-alone block (b). The definition of axes and color are the same as in Figure 4B. **C.** Decoding accuracy in cross-epoch decoding analyses calculated separately for each of the four task phases in the CS-US block (a) and the CS-alone block (b), as in A.

Collectively, these findings suggest that the LEC inhibition abolished the beneficial effect of task demands on ensemble stability and worsened time-dependent drift in the PrL ensemble activity for the learned stimulus associations.

### LEC inhibition decouples PrL firing stability from successful memory expression

Lastly, we investigated how PrL ensemble stability was correlated with CR expression and how it might be affected by LEC inhibition. To this end, we calculated Pearson correlation coefficients between the two firing rate matrices (cells × 4 task phases in each session) constructed from trials in which the rats expressed CRs (CR trials) and those in which they did not (No-CR trials). In aCSF sessions, the correlation coefficients in CR trials were higher than those in No-CR trials (Figure 10), suggesting greater stability of PrL firings when memory was successfully retrieved. Muscimol infusions did not affect this aspect of firing stability in Epoch 3 (two-way mixed ANOVA, CR type×Drug, F_1, 13_ = 1.220, p = 0.289; CR type, F_1, 13_ = 15.204, p = 0.002; Drug, F_1,13_ = 0.901, p = 0.360). In contrast, muscimol infusions eliminated the difference in the stability between CR and No-CR trials in Epoch 4 (CR type×Drug, F_1, 10_ = 5.378, p = 0.043; follow-up paired t-test, aCSF, p = 0.010; muscimol, p = 0.743). These findings suggest that when the rats underwent the stimulus sequence with the disrupted LEC for the second time, behavioral expression of memory was no longer correlated with the stability of stimulus representations in PrL ensembles.

**Figure 10.**
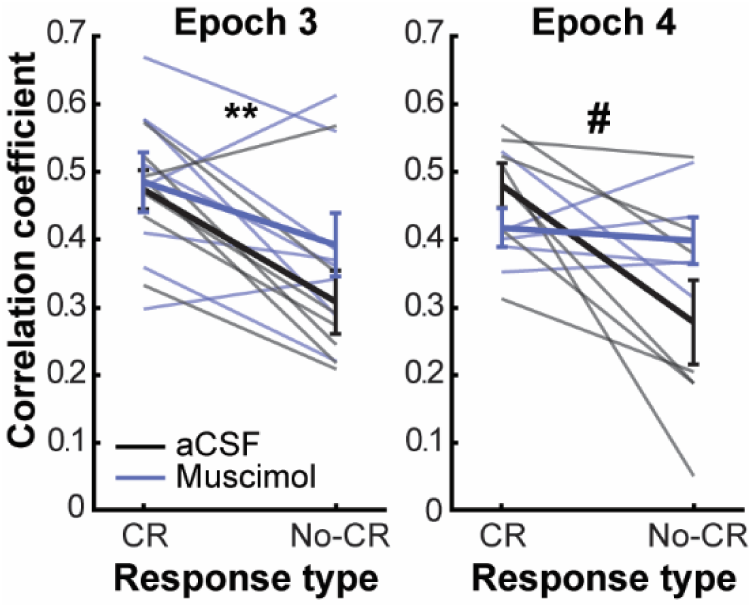
Correlations between the across-trial firing stability and memory expression. The correlation coefficient of ensemble firing rates between two sets of trials with or without CR expression. The analysis was conducted in each session separately and applied only to sessions in which the rats expressed CRs in at least 30% of CS-US trials (aCSF, n = 7 sessions in Epoch 3 and 4; Muscimol, n = 8 sessions in Epoch 3 and 5 sessions in Epoch 4; mean ± s.e.m). ** indicates a significant main effect of Response type (p < 0.01), and # indicates a significant Drug×Response type interaction (p < 0.05) in two-way mixed ANOVA.

## DISCUSSION

Neurons in the LEC represent the conjunction of events, contexts, and their history (Deshmukh and Knierim, 2011; Tsao et al., 2013; Keene et al., 2016; Pilkiw et al., 2017). This unique code, along with its widespread anatomical connectivity (Bota et al., 2015; Nilssen et al., 2019), places the LEC in a strategic position to modulate cortical neural processing during memory retrieval. Here we found that pharmacological inhibition of the LEC destabilized PrL ensemble firings and impaired the reinstatement of neuronal representations of highly familiar experiences.

By comparing multiple single-neuron firings in the PrL across repetitions of constant sensory experiences, we found that LEC inhibition did not affect the encoding of sensory stimuli but instead disrupted their stability over repetitions. Following LEC inhibition, the PrL did not change its magnitude of stimulus-evoked activity (Figure 7Ac, Bc) or its sparsity of stimulus representations (Figure 7Aa, Ba) at the population level. This intact stimulus encoding sharply contrasts with the loss of stimulus-evoked activity in the hippocampus following entorhinal lesion (Ryou et al., 2001) and deafferentation (Woods et al., 2020), suggesting that, unlike the hippocampus, the PrL does not rely on LEC inputs to represent external sensory information (see also, Yu et al., 2021). Instead, LEC inhibition reduced the probability that a consistent set of neurons became activated across repeated experiences (Figure 6C, 7A-C), leading to deficits in two memory-related properties: 1) the reinstatement of the original ensemble state that bridges the temporal gap between the stimulus and the shock (Figure 3C, 4A-C) and 2) the enhanced across-time consistency of ensemble firings by high task demands (Figure 8B). The deficit in the former likely underlies the impaired memory retrieval (Figure 1E) because this particular ensemble state is tightly coupled with the behavioral expression of stimulus-shock associations (Takehara-Nishiuchi et al., 2005; Volle et al., 2016). On the other hand, the latter property is not unique to our behavioral task and was also detected in another subregion in the mPFC (Hyman et al., 2012) and the hippocampus (Kentros et al., 2004; Muzzio et al., 2009; Dupret et al., 2010). These observations suggest that identical external sensory inputs do not guarantee accurate reinstatement of ensemble activity unless animals learn their behavioral relevance (Clopath et al., 2017). The present finding expands this idea by suggesting that the LEC provides a critical signal to stabilize the cortical representations of behaviorally relevant information.

In addition to the major restructuring of ensemble activity during the epoch immediately after LEC inhibition (Epoch 3), the ensemble activity continued to evolve into a new state during a subsequent epoch (Epoch 4; Figure 8C), suggesting exacerbated firing drift over time. Notably, although the new state represented task phases accurately (Figure 9A, B), the across-trial stability of these representations was decoupled from successful memory expression (Figure 10). Such decoupling was not detected in Epoch 3, raising the possibility that “new” learning might occur due to the disrupted memory retrieval by LEC inhibition. Specifically, although the initial training instilled that CS-US contingency changed depending on the temporal context, LEC inhibition may disrupt the retrieval of this contextual information. This would induce extinction learning during the CS-alone block in Epoch 4, leading to the formation of a new ensemble state that is no longer correlated with successful memory retrieval.

Exactly how does the LEC enhance the stability of PrL ensemble activity across repeated relevant experiences? Although the LEC projections terminate on both pyramidal neurons and interneurons in the PrL (Ährlund-Richter et al., 2019), the net effect of LEC inputs appears to be inhibitory: electric stimulation of the LEC induces synaptic inhibition in most PrL pyramidal neurons (Valenti and Grace, 2009). Thus, LEC input may reset the PrL network state by silencing its ongoing activity and maintaining the new state by activating inhibitory-inhibitory connections, as implicated in recent computational models (Mongillo et al., 2018; Kim and Sejnowski, 2021). Of note, parallel to the overall inhibition, LEC stimulation enhances the up states in a small proportion of pyramidal neurons (Valenti and Grace, 2009). Through these effects, the LEC input may depolarize a specific set of PrL pyramidal neurons, thereby enhancing their responses to incoming sensory inputs. Notably, LEC neurons dramatically change their baseline firing rates depending on the history of events experienced in each familiar environment as soon as animals enter the familiar environment (Pilkiw et al., 2017). Such LEC inputs might selectively prime a specific group of PrL pyramidal neurons for upcoming events, thereby supporting their consistent activation across repeated experiences.

One may argue that the LEC’s role in the reinstatement of cortical memory representations is secondary to pattern completion processes in the hippocampus: the LEC dysfunction deprives the hippocampus of sensory inputs necessary for pattern completion and prevents the transfer of the resultant signal to cortical regions. Indeed, some correlational (Solomon et al., 2019; Staresina et al., 2019) and functional evidence (Tanaka et al., 2014) supports the notion that the hippocampus controls the entorhinal cortex during memory retrieval. Several observations, however, point out the importance of LEC local processing for cortical neural reinstatement. Firstly, the neural selectivity for the history of events in a specific environment was detected not only in the deep layers that receive projections from the hippocampus but also in the superficial layers that send projections to the hippocampus (Pilkiw et al., 2017). Thus, the LEC’s unique neural selectivity is not merely a simple copy of hippocampal activity, at least during familiar experiences, suggesting that the LEC can process information independently from the hippocampus (see also, O’Neill et al., 2017). Secondly, the inhibition of other hippocampal output pathways induces qualitatively different changes in frontal firing patterns and behaviour. Specifically, optogenetic inhibition of projections from the ventral hippocampus abolishes the firing selectivity of PrL neurons for goal locations only during the encoding, but not the retrieval phase of the spatial working memory task (Spellman et al., 2015; see also Burton et al., 2009). Furthermore, inhibition of this pathway impairs the learning of new strategies, making mice continue to retrieve the initially learned strategy when the task rule has been changed (Park et al., 2021). In parallel, optogenetic silencing of another output pathway, the ventral subiculum, abolishes the encoding of context-dependent task rules in the orbitofrontal cortex (Wikenheiser et al., 2017). These findings sharply contrast with our observations that LEC inhibition impaired neural reinstatement but not ensemble selectivity for task events (Figures 4, 9). Nonetheless, future investigations with pathway-specific manipulations or hippocampal manipulations are warranted to decipher the precise contribution of the hippocampus and the LEC to cortical neural reinstatement.

Although a failure in neuronal reinstatement is detrimental for memory retrieval, the generation of new ensemble patterns for familiar events may contribute to differentiating the same events that have occurred at different time points (Eichenbaum, 2017; Mau et al., 2020). Accumulating evidence suggests that each brain region is endowed with a mix of stable and changing neural activity patterns across various time scales and magnitude (Clopath et al., 2017; Rule et al., 2019; Mau et al., 2020). Notably, in the LEC, the degree of drift in neural firings is strongly modulated by ongoing experiences: during free foraging in an open environment, LEC ensemble firing patterns change over time scales from seconds to hours; however, such firing drift is markedly suppressed during repeated laps on a figure-eight maze (Tsao et al., 2018). The latter observation is consistent with stable ensemble firing patterns over ∼30 minutes, during which rats received repeated pairings of familiar stimuli in a familiar environment (Pilkiw et al., 2017). These findings suggest that the LEC flexibly switches between an unstable, ever-changing state and a stable, fixed state depending on the familiarity and relevance of present experiences. The former unstable state signals the free-flowing passage of time, supporting single-shot encoding of the temporal order of incidental events in the hippocampus (Rolls and Mills, 2019; Sugar and Moser, 2019). The latter stable state facilitates the reinstatement of the corresponding ensemble code in other brain regions, including the PrL. Therefore, the LEC may provide an essential signal to control the balance between flexibility and stability of cortical neuronal ensembles.

In conclusion, the present findings suggest that the integrity of the LEC is necessary for the reinstatement of fine-scale cortical ensemble representations of learned stimulus associations. Reduced LEC input makes cortical neuronal ensembles enter an unstable state that favors the encoding of present information over the retrieval of past information and impairs memory retrieval. Considering that the LEC is profoundly affected in normal aging and Alzheimer’s disease (Khan et al., 2014; Wisse et al., 2014; Nosheny et al., 2019), we speculate that the imbalance between flexibility and stability in cortical ensemble code might be a mechanism underlying retrieval deficits in the elderly and patients with dementia.

## ACKNOWLEDGEMENTS

This work was supported by CIHR Operating Grant, CFI Leaders Opportunity Fund (K.T.), NSERC graduate fellowship (M.P.), OGS scholarship (J.J.).

## REFERENCES

Ährlund-Richter S, Xuan Y, van Lunteren JA, Kim H, Ortiz C, Pollak Dorocic I, Meletis K, Carlén M (2019) A whole-brain atlas of monosynaptic input targeting four different cell types in the medial prefrontal cortex of the mouse. Nat Neurosci 22:657–668.

Alvarez P, Squire LR (1994) Memory consolidation and the medial temporal lobe: a simple network model. Proc Natl Acad Sci U S A 91:7041–7045.

Bosch SE, Jehee JFM, Fernández G, Doeller CF (2014) Reinstatement of associative memories in early visual cortex is signaled by the hippocampus. J Neurosci Off J Soc Neurosci 34:7493–7500.

Bota M, Sporns O, Swanson LW (2015) Architecture of the cerebral cortical association connectome underlying cognition. Proc Natl Acad Sci U S A 112:E2093–2101.

Burton BG, Hok V, Save E, Poucet B (2009) Lesion of the ventral and intermediate hippocampus abolishes anticipatory activity in the medial prefrontal cortex of the rat. Behav Brain Res 199:222–234.

Chang CC, Lin CJ (2011) LIBSVM:a library for support vector machines. ACM Trans Intell Syst Technol 2:1–27.

Clopath C, Bonhoeffer T, Hübener M, Rose T (2017) Variance and invariance of neuronal long-term representations. Philos Trans R Soc Lond B Biol Sci 372.

Cowansage KK, Shuman T, Dillingham BC, Chang A, Golshani P, Mayford M (2014) Direct reactivation of a coherent neocortical memory of context. Neuron 84:432–441.

Deshmukh SS, Knierim JJ (2011) Representation of non-spatial and spatial information in the lateral entorhinal cortex. Front Behav Neurosci 5:69.

Dupret D, O’Neill J, Pleydell-Bouverie B, Csicsvari J (2010) The reorganization and reactivation of hippocampal maps predict spatial memory performance. Nat Neurosci 13:995–1002.

Eichenbaum H (2000) A cortical-hippocampal system for declarative memory. Nat Rev Neurosci 1:41–50.

Eichenbaum H (2017) Prefrontal-hippocampal interactions in episodic memory. Nat Rev Neurosci 18:547–558.

Frankland PW, Bontempi B (2005) The organization of recent and remote memories. Nat Rev Neurosci 6:119–130.

Goode TD, Tanaka KZ, Sahay A, McHugh TJ (2020) An Integrated Index: Engrams, Place Cells, and Hippocampal Memory. Neuron 107:805–820.

Guo N, Soden ME, Herber C, Kim MT, Besnard A, Lin P, Ma X, Cepko CL, Zweifel LS, Sahay A (2018) Dentate granule cell recruitment of feedforward inhibition governs engram maintenance and remote memory generalization. Nat Med 24:438–449.

Guskjolen A, Kenney JW, de la Parra J, Yeung B-RA, Josselyn SA, Frankland PW (2018) Recovery of “Lost” Infant Memories in Mice. Curr Biol CB 28:2283–2290.e3.

Hainmueller T, Bartos M (2018) Parallel emergence of stable and dynamic memory engrams in the hippocampus. Nature 558:292–296.

Hattori S, Yoon T, Disterhoft JF, Weiss C (2014) Functional reorganization of a prefrontal cortical network mediating consolidation of trace eyeblink conditioning. J Neurosci Off J Soc Neurosci 34:1432–1445.

Horner AJ, Bisby JA, Bush D, Lin W-J, Burgess N (2015) Evidence for holistic episodic recollection via hippocampal pattern completion. Nat Commun 6:7462.

Hyman JM, Ma L, Balaguer-Ballester E, Durstewitz D, Seamans JK (2012) Contextual encoding by ensembles of medial prefrontal cortex neurons. Proc Natl Acad Sci U S A 109:5086– 5091.

Igarashi KM, Lu L, Colgin LL, Moser M-B, Moser EI (2014) Coordination of entorhinal-hippocampal ensemble activity during associative learning. Nature 510:143–147.

Insausti R, Herrero MT, Witter MP (1997) Entorhinal cortex of the rat: cytoarchitectonic subdivisions and the origin and distribution of cortical efferents. Hippocampus 7:146– 183.

Keene CS, Bladon J, McKenzie S, Liu CD, O’Keefe J, Eichenbaum H (2016) Complementary Functional Organization of Neuronal Activity Patterns in the Perirhinal, Lateral Entorhinal, and Medial Entorhinal Cortices. J Neurosci Off J Soc Neurosci 36:3660– 3675.

Kentros CG, Agnihotri NT, Streater S, Hawkins RD, Kandel ER (2004) Increased attention to spatial context increases both place field stability and spatial memory. Neuron 42:283– 295.

Kerr KM, Agster KL, Furtak SC, Burwell RD (2007) Functional neuroanatomy of the parahippocampal region: the lateral and medial entorhinal areas. Hippocampus 17:697– 708.

Khan UA, Liu L, Provenzano FA, Berman DE, Profaci CP, Sloan R, Mayeux R, Duff KE, Small SA (2014) Molecular drivers and cortical spread of lateral entorhinal cortex dysfunction in preclinical Alzheimer’s disease. Nat Neurosci 17:304–311.

Kim JJ, Clark RE, Thompson RF (1995) Hippocampectomy impairs the memory of recently, but not remotely, acquired trace eyeblink conditioned responses. Behav Neurosci 109:195– 203.

Kim R, Sejnowski TJ (2021) Strong inhibitory signaling underlies stable temporal dynamics and working memory in spiking neural networks. Nat Neurosci 24:129–139.

Kloosterman F, Davidson TJ, Gomperts SN, Layton SP, Hale G, Nguyen DP, Wilson MA (2009) Micro-drive array for chronic in vivo recording: drive fabrication. J Vis Exp JoVE.

Knierim JJ, Neunuebel JP, Deshmukh SS (2014) Functional correlates of the lateral and medial entorhinal cortex: objects, path integration and local-global reference frames. Philos Trans R Soc Lond B Biol Sci 369:20130369.

Mankin EA, Sparks FT, Slayyeh B, Sutherland RJ, Leutgeb S, Leutgeb JK (2012) Neuronal code for extended time in the hippocampus. Proc Natl Acad Sci U S A 109:19462–19467.

Martin JH (1991) Autoradiographic estimation of the extent of reversible inactivation produced by microinjection of lidocaine and muscimol in the rat. Neurosci Lett 127:160–164.

Mau W, Hasselmo ME, Cai DJ (2020) The brain in motion: How ensemble fluidity drives memory-updating and flexibility. eLife 9.

McClelland JL, McNaughton BL, O’Reilly RC (1995) Why there are complementary learning systems in the hippocampus and neocortex: insights from the successes and failures of connectionist models of learning and memory. Psychol Rev 102:419–457.

Mongillo G, Rumpel S, Loewenstein Y (2018) Inhibitory connectivity defines the realm of excitatory plasticity. Nat Neurosci 21:1463–1470.

Morrissey MD, Insel N, Takehara-Nishiuchi K (2017) Generalizable knowledge outweighs incidental details in prefrontal ensemble code over time. eLife 6.

Morrissey MD, Maal-Bared G, Brady S, Takehara-Nishiuchi K (2012) Functional dissociation within the entorhinal cortex for memory retrieval of an association between temporally discontiguous stimuli. J Neurosci Off J Soc Neurosci 32:5356–5361.

Muzzio IA, Levita L, Kulkarni J, Monaco J, Kentros C, Stead M, Abbott LF, Kandel ER (2009) Attention enhances the retrieval and stability of visuospatial and olfactory representations in the dorsal hippocampus. PLoS Biol 7:e1000140.

Nilssen ES, Doan TP, Nigro MJ, Ohara S, Witter MP (2019) Neurons and networks in the entorhinal cortex: A reappraisal of the lateral and medial entorhinal subdivisions mediating parallel cortical pathways. Hippocampus 29:1238–1254.

Nosheny RL, Insel PS, Mattsson N, Tosun D, Buckley S, Truran D, Schuff N, Aisen PS, Weiner MW, Alzheimer’s Disease Neuroimaging Initiative (2019) Associations among amyloid status, age, and longitudinal regional brain atrophy in cognitively unimpaired older adults. Neurobiol Aging 82:110–119.

O’Neill J, Boccara CN, Stella F, Schoenenberger P, Csicsvari J (2017) Superficial layers of the medial entorhinal cortex replay independently of the hippocampus. Science 355:184–188.

Park AJ, Harris AZ, Martyniuk KM, Chang C-Y, Abbas AI, Lowes DC, Kellendonk C, Gogos JA, Gordon JA (2021) Reset of hippocampal-prefrontal circuitry facilitates learning. Nature 591:615–619.

Pilkiw M, Insel N, Cui Y, Finney C, Morrissey MD, Takehara-Nishiuchi K (2017) Phasic and tonic neuron ensemble codes for stimulus-environment conjunctions in the lateral entorhinal cortex. eLife 6.

Raposo D, Kaufman MT, Churchland AK (2014) A category-free neural population supports evolving demands during decision-making. Nat Neurosci 17:1784–1792.

Remington ED, Narain D, Hosseini EA, Jazayeri M (2018) Flexible Sensorimotor Computations through Rapid Reconfiguration of Cortical Dynamics. Neuron 98:1005–1019.e5.

Ritchey M, Wing EA, LaBar KS, Cabeza R (2013) Neural similarity between encoding and retrieval is related to memory via hippocampal interactions. Cereb Cortex N Y N 1991 23:2818–2828.

Rolls ET, Mills P (2019) The Generation of Time in the Hippocampal Memory System. Cell Rep 28:1649–1658.e6.

Rule ME, O’Leary T, Harvey CD (2019) Causes and consequences of representational drift. Curr Opin Neurobiol 58:141–147.

Russo AA, Khajeh R, Bittner SR, Perkins SM, Cunningham JP, Abbott LF, Churchland MM (2020) Neural Trajectories in the Supplementary Motor Area and Motor Cortex Exhibit Distinct Geometries, Compatible with Different Classes of Computation. Neuron 107:745–758.e6.

Ryou JW, Cho SY, Kim HT (2001) Lesions of the entorhinal cortex impair acquisition of hippocampal-dependent trace conditioning. Neurobiol Learn Mem 75:121–127.

Schimanski LA, Lipa P, Barnes CA (2013) Tracking the course of hippocampal representations during learning: when is the map required? J Neurosci Off J Soc Neurosci 33:3094–3106.

Solomon EA, Stein JM, Das S, Gorniak R, Sperling MR, Worrell G, Inman CS, Tan RJ, Jobst BC, Rizzuto DS, Kahana MJ (2019) Dynamic Theta Networks in the Human Medial Temporal Lobe Support Episodic Memory. Curr Biol CB 29:1100–1111.e4.

Spellman T, Rigotti M, Ahmari SE, Fusi S, Gogos JA, Gordon JA (2015) Hippocampal-prefrontal input supports spatial encoding in working memory. Nature 522:309–314.

Squire LR (1992) Memory and the hippocampus: a synthesis from findings with rats, monkeys, and humans. Psychol Rev 99:195–231.

Staresina BP, Cooper E, Henson RN (2013) Reversible information flow across the medial temporal lobe: the hippocampus links cortical modules during memory retrieval. J Neurosci Off J Soc Neurosci 33:14184–14192.

Staresina BP, Reber TP, Niediek J, Boström J, Elger CE, Mormann F (2019) Recollection in the human hippocampal-entorhinal cell circuitry. Nat Commun 10:1503.

Sugar J, Moser M-B (2019) Episodic memory: Neuronal codes for what, where, and when. Hippocampus 29:1190–1205.

Suter EE, Weiss C, Disterhoft JF (2019) Differential responsivity of neurons in perirhinal cortex, lateral entorhinal cortex, and dentate gyrus during time-bridging learning. Hippocampus 29:511–526.

Swanson LW, Köhler C (1986) Anatomical evidence for direct projections from the entorhinal area to the entire cortical mantle in the rat. J Neurosci Off J Soc Neurosci 6:3010–3023.

Takehara K, Kawahara S, Kirino Y (2003) Time-dependent reorganization of the brain components underlying memory retention in trace eyeblink conditioning. J Neurosci Off J Soc Neurosci 23:9897–9905.

Takehara-Nishiuchi K (2014) Entorhinal cortex and consolidated memory. Neurosci Res 84:27– 33.

Takehara-Nishiuchi K (2018) The Anatomy and Physiology of Eyeblink Classical Conditioning. Curr Top Behav Neurosci 37:297–323.

Takehara-Nishiuchi K, Kawahara S, Kirino Y (2005) NMDA receptor-dependent processes in the medial prefrontal cortex are important for acquisition and the early stage of consolidation during trace, but not delay eyeblink conditioning. Learn Mem Cold Spring Harb N 12:606–614.

Takehara-Nishiuchi K, Maal-Bared G, Morrissey MD (2011) Increased Entorhinal-Prefrontal Theta Synchronization Parallels Decreased Entorhinal-Hippocampal Theta Synchronization during Learning and Consolidation of Associative Memory. Front Behav Neurosci 5:90.

Takehara-Nishiuchi K, McNaughton BL (2008) Spontaneous changes of neocortical code for associative memory during consolidation. Science 322:960–963.

Takehara-Nishiuchi K, Morrissey MD, Pilkiw M (2020) Prefrontal Neural Ensembles Develop Selective Code for Stimulus Associations within Minutes of Novel Experiences. J Neurosci Off J Soc Neurosci 40:8355–8366.

Takehara-Nishiuchi K, Nakao K, Kawahara S, Matsuki N, Kirino Y (2006) Systems consolidation requires postlearning activation of NMDA receptors in the medial prefrontal cortex in trace eyeblink conditioning. J Neurosci Off J Soc Neurosci 26:5049– 5058.

Tanaka KZ, Pevzner A, Hamidi AB, Nakazawa Y, Graham J, Wiltgen BJ (2014) Cortical representations are reinstated by the hippocampus during memory retrieval. Neuron 84:347–354.

Taxidis J, Pnevmatikakis EA, Dorian CC, Mylavarapu AL, Arora JS, Samadian KD, Hoffberg EA, Golshani P (2020) Differential Emergence and Stability of Sensory and Temporal Representations in Context-Specific Hippocampal Sequences. Neuron 108:984–998.e9.

Teyler TJ, DiScenna P (1986) The hippocampal memory indexing theory. Behav Neurosci 100:147–154.

Teyler TJ, Rudy JW (2007) The hippocampal indexing theory and episodic memory: updating the index. Hippocampus 17:1158–1169.

Tsao A, Moser M-B, Moser EI (2013) Traces of experience in the lateral entorhinal cortex. Curr Biol CB 23:399–405.

Tsao A, Sugar J, Lu L, Wang C, Knierim JJ, Moser M-B, Moser EI (2018) Integrating time from experience in the lateral entorhinal cortex. Nature 561:57–62.

Valenti O, Grace AA (2009) Entorhinal cortex inhibits medial prefrontal cortex and modulates the activity states of electrophysiologically characterized pyramidal neurons in vivo. Cereb Cortex N Y N 1991 19:658–674.

Volle J, Yu X, Sun H, Tanninen SE, Insel N, Takehara-Nishiuchi K (2016) Enhancing Prefrontal Neuron Activity Enables Associative Learning of Temporally Disparate Events. Cell Rep 15:2400–2410.

Wikenheiser AM, Marrero-Garcia Y, Schoenbaum G (2017) Suppression of Ventral Hippocampal Output Impairs Integrated Orbitofrontal Encoding of Task Structure. Neuron 95:1197–1207.e3.

Wilson MA, McNaughton BL (1993) Dynamics of the hippocampal ensemble code for space. Science 261:1055–1058.

Wisse LEM, Biessels GJ, Heringa SM, Kuijf HJ, Koek DHL, Luijten PR, Geerlings MI, Utrecht Vascular Cognitive Impairment (VCI) Study Group (2014) Hippocampal subfield volumes at 7T in early Alzheimer’s disease and normal aging. Neurobiol Aging 35:2039– 2045.

Woodruff-Pak DS, Disterhoft JF (2008) Where is the trace in trace conditioning? Trends Neurosci 31:105–112.

Woods NI, Stefanini F, Apodaca-Montano DL, Tan IMC, Biane JS, Kheirbek MA (2020) The Dentate Gyrus Classifies Cortical Representations of Learned Stimuli. Neuron 107:173–184.e6.

Xing B, Morrissey MD, Takehara-Nishiuchi K (2020) Distributed representations of temporal stimulus associations across regular-firing and fast-spiking neurons in rat medial prefrontal cortex. J Neurophysiol 123:439–450.

Xu W, Wilson DA (2012) Odor-evoked activity in the mouse lateral entorhinal cortex. Neuroscience 223:12–20.

Young BJ, Otto T, Fox GD, Eichenbaum H (1997) Memory representation within the parahippocampal region. J Neurosci Off J Soc Neurosci 17:5183–5195.

Yu XT, Yu J, Choi A, Takehara-Nishiuchi K (2021) Lateral entorhinal cortex supports the development of prefrontal network activity that bridges temporally discontiguous stimuli. Hippocampus, *in press*.

Ziv Y, Burns LD, Cocker ED, Hamel EO, Ghosh KK, Kitch LJ, El Gamal A, Schnitzer MJ (2013) Long-term dynamics of CA1 hippocampal place codes. Nat Neurosci 16:264–266.

